# Lymphotoxin*-*β Promotes Bone Colonization and Osteolytic Outgrowth of Indolent Bone Metastatic cells of Breast Cancer

**DOI:** 10.1101/2023.08.15.553179

**Authors:** Xuxiang Wang, Tengjiang Zhang, Bingxin Zheng, Youxue Lu, Yong Liang, Guoyuan Xu, Luyang Zhao, Yuwei Tao, Qianhui Song, Huiwen You, Haitian Hu, Xuan Li, Keyong Sun, Tianqi Li, Zian Zhang, Jianbin Wang, Xun Lan, Deng Pan, Yang-Xin Fu, Bin Yue, Hanqiu Zheng

## Abstract

Bone metastatic relapse is a lethal consequence of breast cancer, occurring years after initial diagnosis. By analyzing single-cell transcriptomes of bone-seeding tumor cells and *in vivo* barcoded cDNA library screening, LTβ (lymphotoxin-β) is identified as a key factor highly expressed in early-stage bone metastatic cells, associated with poor bone metastasis-free survival, and capable of promoting dormancy reactivation in multiple breast cancer models. Mechanistically, tumor-derived LTβ activates NF-κB2 signaling in osteoblasts to express CCL2/5, facilitating tumor cell seeding and accelerating osteoclastogenesis. Both processes contribute to the reactivation of dormancy and metastatic outgrowth. Blocking LTβ signaling with a decoy receptor significantly suppressed bone colonization and metastatic progression, whereas clinical sample analysis revealed significantly higher LTβ expression in bone metastases than in primary tumors. Our findings highlight LTβ as a bone niche-induced factor that promotes tumor cell seeding and dormancy reactivation, underscoring its potential as a therapeutic target for preventing bone metastatic relapse in patients with breast cancer.

## INTRODUCTION

Bone metastasis is a common and devastating complication of breast cancer, with over 70% of patients eventually developing it (Esposito et al., 2018). Bone metastasis is associated with severe symptoms, such as bone pain, fractures, cachexia, and hypercalcemia, which can be life-threatening (Waning et al., 2015; Weilbaecher et al., 2011). Additionally, cancer cells within the bone niche could act as “cancer seeds”, spreading to other organs and leading to widespread metastasis (Zhang et al., 2021). Even after successful surgical removal of the primary tumor, dormant cells may remain in the body, contributing to bone relapse. This phenomenon suggests the very low efficiency of bone metastasis colonization and highlights the importance of understanding the molecular mechanisms of this early bone-seeding phase (Elkholi et al., 2022; Uhr and Pantel, 2011).

Studies have highlighted the importance of interactions between tumor cells and the microenvironment, which mainly consists of osteoblasts (Wang et al., 2015; Zheng et al., 2017), osteoclasts (Esposito *et al*., 2018; Lu et al., 2011) and perivascular cells (Ghajar et al., 2013), in the development of bone metastasis. Many critical molecules have been identified to mediate these tumor-stromal interactions during bone metastasis progression, including parathyroid hormone-related peptide (PTHrP) (Yin et al., 1999) which induces the osteolytic cycle (Coleman, 2001; Roodman, 2004) and other factors, such as vascular cell adhesion protein 1 (VCAM-1) (Lu *et al*., 2011), integrin β3 (Ross et al., 2017), and Jagged1-Notch signaling (Sethi et al., 2011), which further enhance this cycle. The cooperation of these factors leads to a vicious cycle of bone degradation and metastatic outgrowth (Satcher and Zhang, 2021). Among the multi-step metastatic cascades, colonization (early seeding event) is the least efficient but potentially the most critical step for successful bone metastasis outgrowth (Massague and Obenauf, 2016). In contrast to the well-studied signals in late-stage osteolytic bone metastasis, the molecular signals in early-stage bone colonization are not well understood.

Understanding the mechanisms controlling tumor cell dormancy and reactivation is crucial for developing therapeutic strategies against cancer (Hu et al., 2023; Malladi et al., 2016; Sosa et al., 2014). A few studies have reported different microenvironmental changes that induce the reawakening of indolent metastatic cells or therapy resistance at multiple metastatic sites. For instance, perivascular-derived thrombospondin-1 induces breast cancer dormancy in lung metastases (Ghajar *et al*., 2013). Astrocyte-deposited laminin-211 induces disseminated breast tumor cell dormancy in the brain (Dai et al., 2022). In the bone marrow microenvironment, study suggests that mesenchymal stem cells (MSCs) promote metastatic breast cancer cells to enter dormancy (Nobre et al., 2021). Another study revealed a role of IL-6 cytokine leukemia inhibitory factor (LIF) helps to maintain cancer dormancy in the bone (Johnson et al., 2016). Dormant tumor cells in the bone may be considered as trapped in the “extended seeding phase” before the eventual outgrowth. There might be common signaling molecules mediating successful bone metastatic seeding and the outbreak of indolent bone metastatic cells. Thus, molecular mediators that facilitate the bone colonization might also be critical to the re-activation of indolent tumor cells in the bone.

Many genes involved in bone metastasis have been identified through gene expression profiling of either *in vivo* selected bone metastasis-prone cells or *in vitro* generated single colonies intrinsically possessing bone metastasis-promoting ability(Kang et al., 2003; Lelekakis et al., 1999; Lu *et al*., 2011; Minn et al., 2005b; Sethi *et al*., 2011). The rationale behind this is that cancer cell lines are heterogenous, and a rare subset of cancer cells possess the ability to establish bone metastasis when they are in the bone microenvironment. However, the bone microenvironment derived cues could affect tumor state and induce critical gene expression, which can contribute to bone colonization and metastasis. These critical signals could not be identified by gene profiling after *in vitro* cell culture. Therefore, it is important to conduct true gene profiling of cancer cells at its *in situ* and *in vivo* state during early-seeding phase. However, because of the very limited number of colonization tumor cells could be recovered, the exact transcriptome profiles of cancer cells during colonization have been missing. Single-cell sequencing (scRNA-seq) technology has revolutionized the study of population dynamics in multiple biological processes(Blake et al., 2015; Gao et al., 2018; Li et al., 2017; Navin et al., 2011). With its ability to determine the transcriptome of a single cell, scRNA-seq technology provided us a potential tool to reveal the early-seeding signaling during bone metastasis.

In the current study, we conducted scRNA-seq analysis on tumor cells obtained from early-stage bone colonization in experimental bone metastasis models and identified potential factors that promote tumor cell survival during the seeding phase. Using *in vivo* mini-cDNA library screening and validation, we discovered that lymphotoxin-β (LTβ), a member of the tumor necrosis factor (TNF) superfamily, enhances breast cancer bone colonization and the reactivation of indolent bone metastatic cells.

## RESULTS

### Determination of the early bone-seeding phase

The 4T1 cell series consists of mouse mammary tumor cell lines that exhibit varying potential for bone metastasis. While 4T1 is known to strongly develop lung metastases, it only weakly metastasizes to bone (Aslakson and Miller, 1992). In contrast, 4T1.2, a single-cell clone derived from 4T1, has a strong propensity to metastasize to bone (Lelekakis *et al*., 1999). Both cell lines were labeled with GFP and *Firefly-luciferase* for live animal *in vivo* bioluminescence imaging (BLI). Our experimental bone metastasis assay confirmed that 4T1.2 generated strong bone metastasis in the majority of mice, while less than 40% of mice injected with 4T1 developed bone metastasis (Supplementary Fig. S1A). Importantly, one day post intracardiac (IC) injection, the BLI signal dropped sharply to the lowest point in both cell lines, and then stabilized at that level before picking up the signal at around 4-5 days (Supplementary Fig. S1B). In both 4T1 and 4T1.2 cell models, osteolytic bone degradation was only apparent at the late stage through μCT imaging and by tartrate-resistant acid phosphatase-positive (TRAP^+^) osteoclasts staining (Supplementary Fig. S1C and data not shown). Similar BLI signal changes could be observed in other bone metastasis cell lines (data not shown).

Based on these observations, we determined that the seeding phase of bone metastasis lasts about a week post injection, and it is safe to recover cancer cells from the bones of IC-injected mice at Day 4 to capture the early bone colonization phase. To investigate the critical mediators of bone colonization, we employed the SMART-seq2 method for single-cell RNA sequencing analysis to examine early bone seeding tumor cells (Picelli et al., 2014) with experimental design illustrated in Fig. 1A. Although this protocol has low throughput, it allows for the detection of a large number of genes in each cell, particularly those with low abundance (Wang et al., 2021). Cells labeled with *Firefly-luciferase* and mCherry were IC injected into mice. At different days post injection, a group of mice was sacrificed to recover bone metastatic cancer cells from the hindlimbs. mCherry^+^ tumor cells were isolated using fluorescence-activated cell sorting (FACS). Single-cell cDNA amplification and next-generation sequencing (NGS) library construction were performed. *In vitro* cultured cells and mammary fat-pad injected primary tumor cells were isolated to generate scRNA-seq results as additional controls. Candidate genes were chosen based on scRNA-seq data, and a cDNA overexpression library was constructed for *in vivo* bone metastasis screening. Further mechanistic studies were conducted to reveal the detailed tumor-stromal crosstalk that promotes bone metastasis (Fig. 1A).

**Figure 1.**
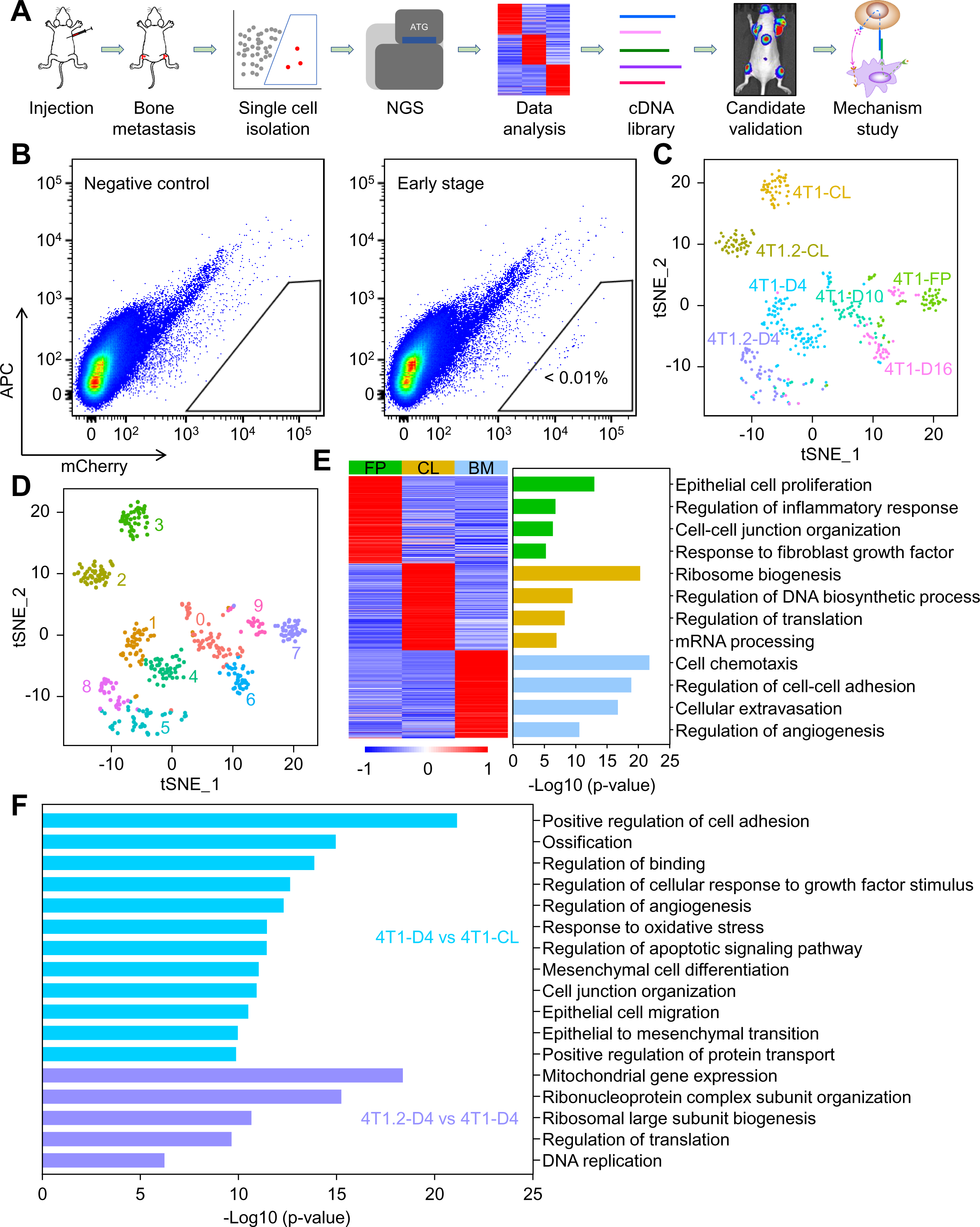
Single Cell RNA-Seq Analysis of Cancer Cells from Different Stages of Bone Metastasis. **(A)** Schematic representation of the experiment design. mCherry-labeled cancer cells were IC injected into female BALB/c mice for bone metastasis formation. Cells were collected by FACS sorting at different time points post injection. ScRNA-Seq and bioinformatic analysis were performed to identify bone metastatic seeding-promoting genes. Candidate genes were further validated by mini-cDNA library screen in bone metastasis assay *in vivo*. The top candidate genes were then selected for functional and mechanistic studies. **(B)** Representative FACS-plot of control mice and mCherry^+^ cancer cell injected mice of early-stage bone metastasis were displayed. Female mice were IC injected with PBS or mCherry^+^ tumor cells to generate bone metastasis. Four days post injection, mice were sacrificed for hindlimb bones. Bones were minced and digested for cell suspension. mCherry^+^ cancer cells were isolated by FACS-sorting. (**C**) ScRNA sequencing was performed with cells from *in vitro* culture, primary tumors from fat-pad injection, and different stages of bone metastasis. t-SNE plot of all these cancer cells were presented. Tumor cells from different sources were color-coded. (**D**) Similar as in **C**, t-SNE analysis of all cells were presented. Different cell clusters were color-coded. (**E**) Heatmap and GO enriched terms of 4T1 cells from fat pad injection (FP), *in vitro* cultured cells (CL), and bone metastasis (BM). (**F**) GO analysis of enriched terms in 4T1-D4 cells *v.s.* 4T1-CL cells and 4T1.2-D4 cells *v.s.* 4T1-D4 cells. **See also Supplementary Figure S1.**

### Single cell transcriptome profiling of tumor cells in different phases of bone metastasis

The low percentage of mCherry^+^ tumor cells in the bone marrow day 4 post injection suggests that metastatic seeding is inefficient (Fig. 1B) (Lambert et al., 2017). The isolated tumor cells were directly used for scRNA-seq without additional *in vitro* culture to preserve their native transcriptomes. We acquired high-quality transcriptome profiles of 424 cells (Supplementary Table S1). In short, we acquired a full set of scRNA-seq data of 4T1 cells from mammary fat pad injection (FP), *in vitro* cultured cells (CL), Day 4 (early seeding phase, D4), Day 10 (progression phase, D10), Day 16 (late-stage bone metastasis phase, D16) post IC injection. Additionally, we also acquired a partial set of scRNA-seq data of 4T1.2 cells from CL and D4. A t-distributed stochastic neighbor embedding (t-SNE) analysis identified ten distinct clusters (Figs. 1C&D), with most cells homogeneously expressing *Firefly-luciferase* and *mCherry*, confirming their tumor cell identities (Supplementary Fig. S1D). The authenticity of the isolated tumor cells was additionally confirmed through the expression of *Keratin 8* (*Krt8*), a marker for epithelial origin. *Vcam-1*, a previously reported bone metastasis-promoting gene, was highly expressed in tumor cells isolated from bone marrow, further confirming our scRNA-seq results (Supplementary Fig. S1D)

In the t-SNE analysis, 4T1-CL and 4T1.2-CL cell populations were clearly distinguishable from other cell clusters, while the 4T1-FP cells formed a distinct population. Additionally, tumor cells isolated from the bone marrow were more closely related and formed clusters based on the number of days post injection. To further investigate bone metastasis progression, we used cells from FP, CL, D4, D10, and D16 from the 4T1 scRNA-seq dataset, which provided a complete snapshot of the metastatic cascade (Massague and Obenauf, 2016). Gene Ontology (GO) analysis of cells from bone metastases (BM) showed enrichment in cell chemotaxis, cell-cell adhesion, cellular extravasation, and regulation of angiogenesis, highlighting the potential importance of tumor-stromal interactions in bone metastasis (Fig. 1E). To examine the cell transition trajectory, we conducted pseudotime analysis (Reid and Wernisch, 2016), which suggested that 4T1-D4 cells potentially evolved through 4T1-D10 status before becoming 4T1-D16, the osteolytic bone metastasis phase. Notably, CL and FP cells were significantly different from early-stage (D4) or late-stage (D16) bone metastatic cells, highlighting the importance of the bone microenvironment in programming the transcriptome of bone-seeding cells (Supplementary Figs. S1E&F).

Bone metastasis colonization is strongly influenced by tumor-stroma interactions (Massague and Obenauf, 2016; Wang *et al*., 2015; Zheng et al., 2016). We analyzed early “seeding” signals by comparing the gene expression profiles of early-stage bone metastasis (4T1-D4) to *in vitro* cultured cells (4T1-CL), instead of cells from primary tumors (FP). This comparison helps to eliminate the potential noisy signal due to the microenvironment effect in FP group. GO analysis showed that signaling pathways involved in tumor-stromal interactions, such as positive regulation of cell adhesion, cell junction organization, and cellular response to growth factor stimulus, were enriched in the early-stage bone metastasis group (Fig. 1F). As the 4T1.2 cell line has a stronger bone metastasis potential than 4T1, we also compared their gene expression profiles during the early-seeding phase (4T1.2-D4 vs. 4T1-D4) (Fig. 1F). GO terms enriched in 4T1.2-D4 cells were related to cell proliferation pathways such as ribonucleoprotein complex subunit organization, regulation of translation, and DNA replication, suggesting a higher proliferation rate of 4T1.2 cells in the bone microenvironment.

EMT and MET are cellular processes in which cells undergo transformation from epithelial to mesenchymal or from mesenchymal to epithelial, respectively. These processes have been widely studied for their roles in cancer metastasis (Jolly et al., 2017; Thiery et al., 2009). The EMT status of cells during bone metastasis was analyzed at single-cell resolution by using single-cell transcriptomes. Cells from the FP and CL expressed high levels of epithelial cell markers, whereas D4 cells were mesenchymal-like (Supplementary Figs. S1G&H). Interestingly, the epithelial scores of the individual cells varied greatly, indicating EMT plasticity during cancer cell colonization. As tumor cells progressed to generate overt bone metastasis (D10 and D16 cells), a large portion of cells gradually regained their epithelial properties (Supplementary Fig. S1G). This data suggests a potential functional involvement of EMT and MET in bone metastasis progression. Further functional testing could be carried out in the future.

Here, we present a comprehensive global gene expression profile of cells at various stages of bone metastasis, achieved through scRNA-seq analysis. The enrichment of tumor-stromal interaction signals highlights the importance of tumor microenvironment in determining bone metastasis progression.

### Barcoded cDNA library screening identified candidate genes for promoting bone metastasis

A barcoded cDNA library was designed for the *in vivo* screening of genes promoting bone metastasis based on scRNA-seq data. Candidate genes were selected from highly expressed genes by two comparisons illustrated in Fig. 1F, resulting in 84 successfully cloned genes with three unique 10-bp barcodes each (Fig. 2A and Supplementary Table S2). The representative gene expression patterns in t-SNE plot (top five genes in the combined list in Supplementary Table 2) were shown in Supplementary Fig. S2A.

**Figure 2.**
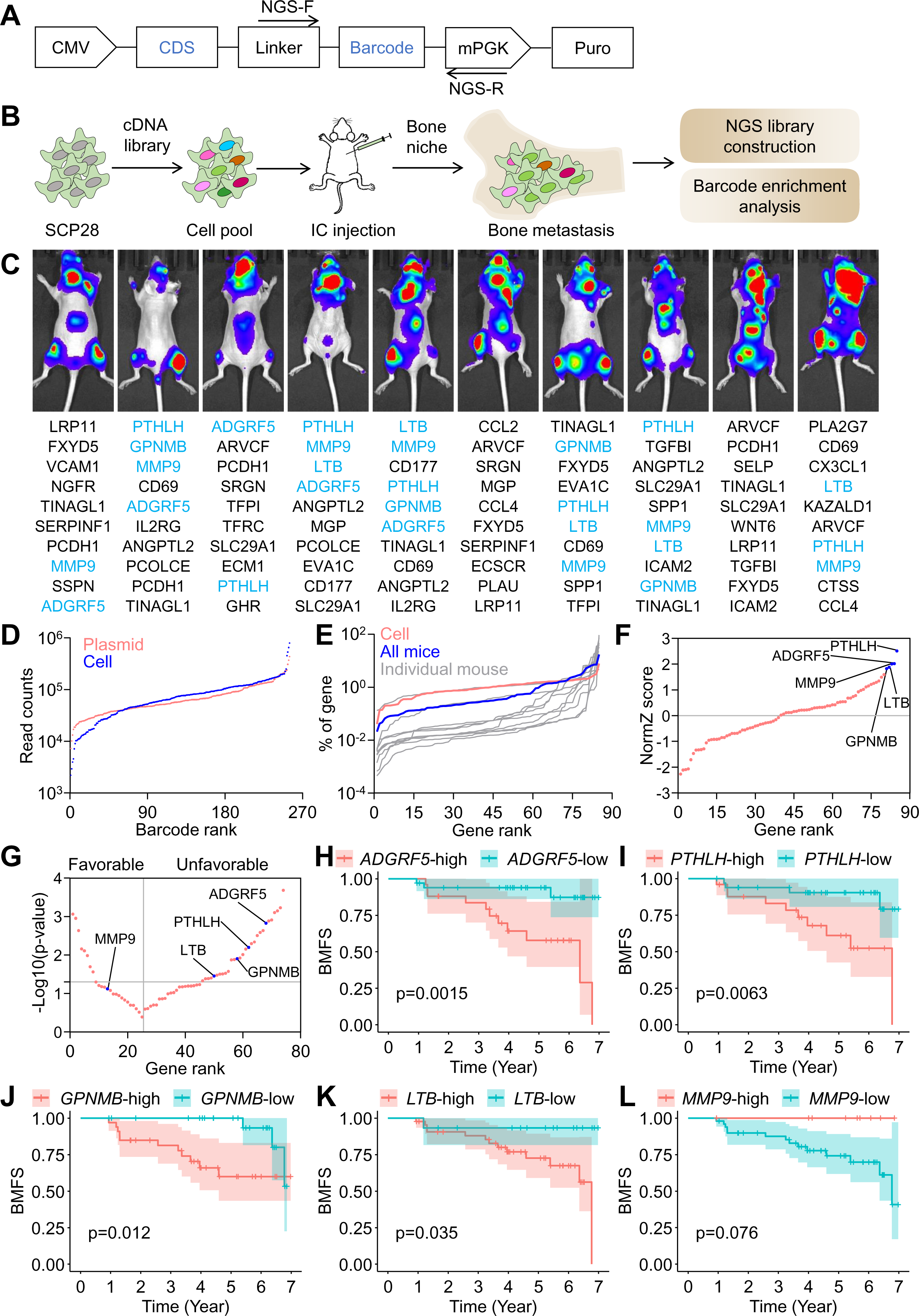
A Barcoded Mini-cDNA Library Screen Identifies Top Candidate Genes for Bone Colonization and Metastatic Progression. (**A**) Schematic diagram of expression vector design for barcoded cDNA library. CMV, human cytomegalovirus (CMV) immediate-early promoter; CDS, coding sequence for each candidate gene; Linker, a DNA sequence gapping the CDS and barcode sequence; Barcode, a 10-bp DNA sequence; mPGK, murine phosphoglycerate kinase (PGK) promoter; Puro, puromycin resistance gene. NGS-F and NGS-R, primer for amplification of barcode sequence, detailed DNA sequences were included in Supplementary Table S4. (**B**) Schematic diagram of *in vivo* mini-cDNA library screen. SCP28 cells were infected with mini-cDNA library. Cells were then IC injected into female nude mice to generate bone metastasis. Six weeks after IC injection, mice were sacrificed and hindlimb bones were collected and minced for direct NGS library construction. NGS library was sequenced for barcodes (and genes) enrichment analysis. (**C**) BLI images of all mice at the experimental endpoints. The top enriched genes in each mouse were presented. Note that the light blue color-labeled genes were top candidate genes that were repeatedly enriched in these mice. (**D**) Ranked barcode abundance in the plasmid library and in cancer cells transduced with lentiviral library as determined by raw read counts from NGS. (**E**) Ranked gene abundance in cancer cells transduced with lentiviral library, in metastases from individual mouse or from all mice as determined by the percentage of each barcode from NGS. (**F**) Ranked gene plot for bone metastasis enriched genes according to their enrichment NormZ score. Note that the most enriched genes are *PTHLH*, *ADGRF5*, *MMP9*, *LTB*, and *GPNMB*. (**G**) Gene ranked by the p-values of Kaplan-Meier curve of BMFS in breast cancer patient data set, stratified by the individual gene expression (Minn *et al*., 2005a). Genes on the left were those whose higher expressions were associated with better patient survival; while genes on the right were those whose higher expressions were associated with worse patient survival. (**H-L**) Kaplan-Meier plots of BMFS in a breast cancer patient data set, stratified by the expression of each candidate gene (*ADGRF5, PTHLH, GPNMB, LTB*, and *MMP9*) (Minn *et al*., 2005a). **See also Supplementary Figure S2.**

An *in vivo* screen was performed as illustrated in Fig. 2B. To make sure the identified genes were functionally important in human cancer, we chose SCP28, a single cell-derived progeny from human MDA-MB-231 breast cancer cell line with moderate bone metastasis potential, as a model cell line for this screen (Kang *et al*., 2003; Minn *et al*., 2005b). A lentivirus-based cDNA library was used to infect SCP28 cells, and *in vivo* bone metastasis screening was performed in 6-week-old female nude mice with a coverage rate of over 10,000× for each barcoded clone. BLI was used to confirm the bone metastasis burden, and the hindlimbs were dissected for bone tissue collection. DNA was extracted from the bone tissues of individual mice, and a next-generation sequencing (NGS) library was generated for barcode detection and enrichment analysis (Fig. 2B).

The initial barcode representation was confirmed to be evenly distributed in the plasmid library and in the infected cells (Fig. 2D). While the majority of genes were depleted after *in vivo* selection, a small number of genes were significantly enriched in bone metastasis samples repeatedly in multiple independent mouse samples (Figs. 2C, E). Gene enrichment analysis showed significant enrichment of *PTHLH*, *ADGRF5*, *MMP9*, *LTB*, and *GPNMB* in bone metastasis samples compared with *in vitro* cultured cells (Fig. 2F) (Colic et al., 2019). As expected, these five genes were highly expressed in bone metastasis samples, but not in cell lines or primary tumor cells, or highly expressed in 4T1.2-D4, but not in 4T1-D4 (Supplementary Fig. S2B). We then assessed the association of these genes with bone metastasis-free survival (BMFS) in the Minn et al. dataset and ranked them based on their P-values in their respective Kaplan-Meier plots (Fig. 2G) (Gyorffy, 2021; Minn et al., 2005a). Among these five genes, higher mRNA levels of *ADGRF5*, *PTHLH*, *GPNMB*, and *LTB* were correlated with shorter BMFS in patients with breast cancer (Figs. 2H-K). Conversely, higher mRNA levels of *MMP9* correlated with longer BMFS (Fig. 2L). We further analyzed the relapse-free survival (RFS) in lymph node-positive breast cancer patients using an independent patient dataset (Lanczky and Gyorffy, 2021). Of these five candidates, four were correlated with worse RFS, whereas *PTHLH* was not (Supplementary Figs. S2C-H).

Previous studies have reported the involvement of osteoactivin (encoded by *GPNMB*) in promoting the progression of bone metastasis, which supports our findings (Maric et al., 2013; Rose et al., 2007; Rose et al., 2017). One study suggested the possible involvement of ADGRF5 in bone metastasis (Tang et al., 2013). On the contrary, the role of LTβ (encoded by *LTB*) in this process remains unknown, further functional validation is needed to confirm the roles of LTβ and ADGRF5 in bone colonization and metastasis.

### LTβ enhances bone metastasis progression

To confirm the potential role of LTβ and ADGRF5 in bone metastasis, we first overexpressed them in SCP28 cells (Supplementary Figs. S3A&B). Overexpression of either LTβ or ADGRF5 had no impact on their cell proliferation rates *in vitro* (Supplementary Fig. S3C). These cells were then IC injected into nude mice for bone metastasis analysis. LTβ-OE significantly increased bone metastasis burden, as quantified by BLI, whereas ADGRF5-OE had no effect on bone metastasis progression (Supplementary Fig. S3D). Histological analysis further demonstrated that LTβ-OE increased the osteolytic lesion area and the number of TRAP^+^ osteoclasts, whereas ADGRF5-OE had much less effects (Supplementary Fig. S3E-G).

To complement the overexpression study, we used short hairpin RNAs (shRNAs) to knock down (KD) LTβ and ADGRF5 individually in SCP28 cells (Supplementary Figs. S3H&I). KD of LTβ significantly decreased the metastatic burden and extended the BMFS of mice (Figs. 3A&B). μCT imaging revealed that severe bone destruction occurred in the scramble shRNA control group, while bone integrity was well preserved in the LTβ KD group (Fig. 3C and Supplementary Videos 1-3). Consistent with the BLI and μCT results, LTβ KD also decreased the osteolytic tumor area in the bone and reduced osteoclast activation within metastatic areas (H&E and TRAP staining in Figs. 3C-E). Interestingly, the KD of ADGRF5 did not reduce the bone metastasis burden or extend the survival time compared to that in the control group (Figs. 3A-C). However, μCT imaging revealed that ADGRF5 KD protected bone from degradation and reduced the number of mature TRAP^+^ osteoclasts to some extent, although its effect is much weaker compared to LTβ KD group (Figs. 3C-E).

**Figure 3.**
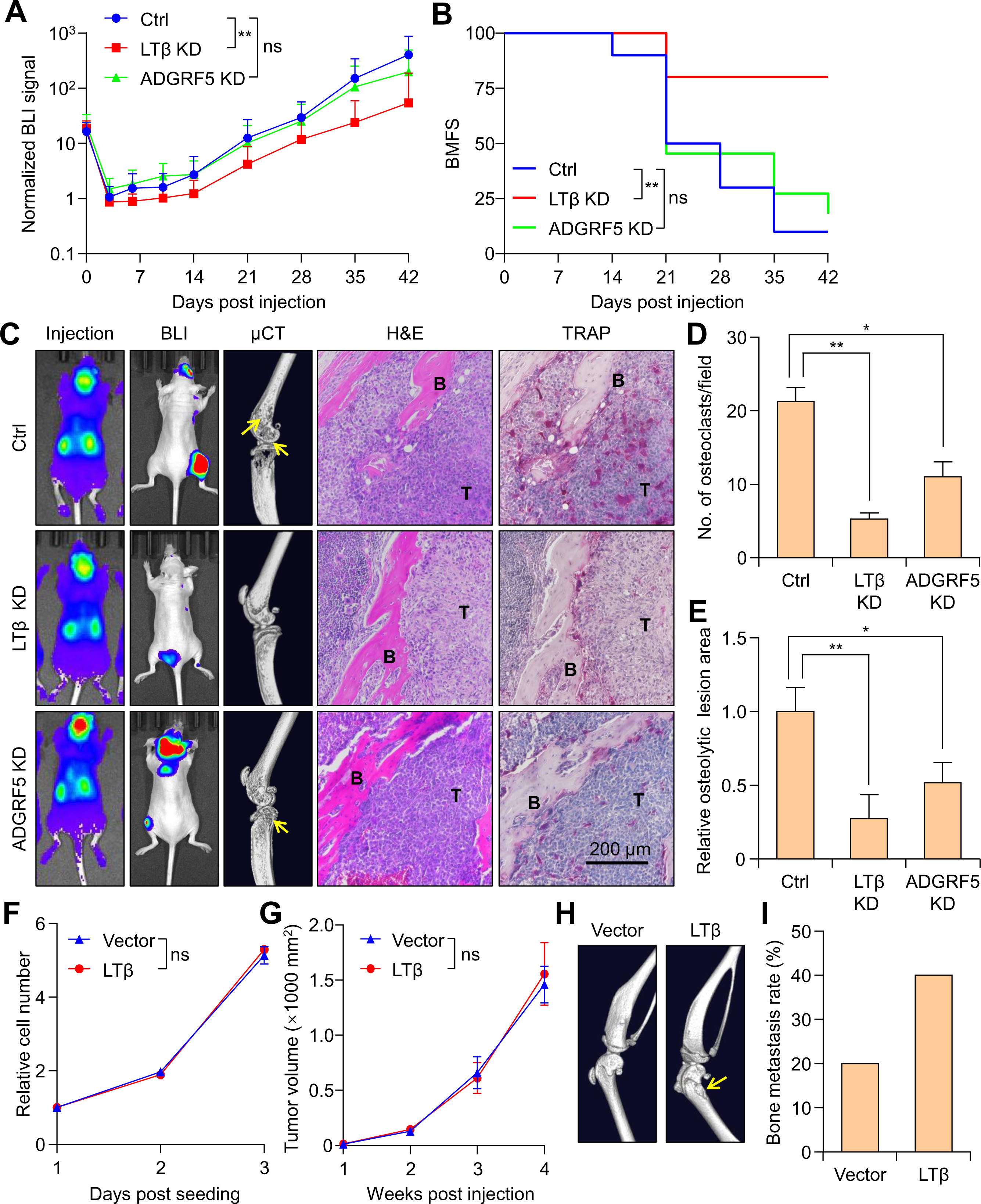
LTβ Emerges as the Top Candidate for Promoting Bone Metastasis. (**A**) SCP28 cells were KD by shRNAs against *LTB* or *ADGRF5*. Scramble shRNA was utilized as negative control. 10^5^ tumor cells were IC injected into 6 weeks old female nude mice. Quantification of bone metastasis burden each week based on BLI imaging. n = 10 mice for Control group, n = 10 mice for LTβ KD group, and n = 11 mice for ADGRF5 KD group. Data presented as mean ± SD. ns: p > 0.05, **p < 0.01 by two-way repeated measures ANOVA. (**B**) Kaplan-Meier curve of mice bearing bone metastasis in experiment performed in **A**. n = 10 mice for Control group, n = 10 mice for LTβ KD group, and n = 11 mice for ADGRF5 KD group. ns: p > 0.05, **p < 0.01 by log-rank test. (**C**) Representative BLI (Day 0 and Day 42), μCT, H&E, and TRAP staining images (Day 42) from mice in experiment performed in **A**. Arrows indicate the osteolytic bone lesion areas. B, bone tissue area; T, tumor area. Scale bar = 200 μm. (**D**) Quantification of TRAP^+^ osteoclasts from decalcified histological bone sections of hindlimbs from mice in **C**, n = 4 per group. Data presented as mean ± SEM. *p < 0.05, **p < 0.01 by unpaired t-test. (**E**) Quantification of osteolytic lesions based on μCT images from **C**. n = 10 mice for control group, n =10 mice for LTβ KD group, and n = 11 mice for ADGRF5 KD group. Data presented as mean ± SEM. *p < 0.05, **p < 0.01 by unpaired t-test. (**F**) 10^3^ 4T1 cells with-Vector control or LTβ-OE were seeded into 96 well plate for culture. Relative cell number was monitored by *in vitro* luciferase assay. n = 4 for each group. Data presented as mean ± SD. “ns” means not statistically different by two-way repeated measures ANOVA. (**G**) 4T1 cells with-Vector control or LTβ-OE were injected in the mammary fat pads of 6-week-old female BALB/c mice. Tumor volume was measured weekly. n = 10 mice per group. (**H**) Representative μCT images of the bones of experimental mice in **G** at experimental endpoint. Arrows indicate the osteolytic bone lesion areas. (**I**) Quantification of the percentage of mice which developed spontaneous bone metastasis based on μCT imaging. **See also Supplementary Figure S3 and Supplementary Videos S1-3.**

We tested whether LTβ could affect primary tumor progression. To this end, SCP28-Vector or SCP28-LTβ cells were FP injected into female athymic nude mice and primary tumor growth was recorded. There was no significant difference between the two groups (Supplementary Fig. S3J). We then utilized the 4T1 cell model and generated either Vector control or LTβ-OE cell line. LTβ-OE did not increase cell proliferation rate *in vitro* (Fig. 3F). These cells were FP injected into BALB/c syngeneic mice. Again, the expression of LTβ did not accelerate primary tumor growth (Fig. 3G). At the experimental endpoint, we sacrificed the mice for osteolytic bone metastasis detection by μCT Imaging. Significantly more mice bearing LTβ-OE cells generated overt osteolytic bone metastasis (Figs. 3H&I). These results suggest that LTβ facilitates bone metastasis progression in spontaneous tumor model without affecting primary tumor growth.

We investigated the significant role of LTβ in bone metastasis through both overexpression and knockdown strategies, utilizing intracardiac injection and spontaneous, immune-competent mouse models. Our findings demonstrate that LTβ promotes bone metastasis progression while leaving primary tumor growth unaffected.

### LTβ reactivates indolent metastatic cancer cells to generate late-stage, osteolytic bone metastasis

The high expression level of LTβ during early-stage bone colonization suggests that it may play a role in promoting the reactivation of dormant tumor cells. To investigate this hypothesis, we employed an MDA-MB-231 derivative cell line named PD2R. PD2R is characterized as an indolent bone metastatic cell line that lacks the ability to develop bone metastasis (Lu *et al*., 2011). Indeed, unlike SCP28 cells, when IC injected into athymic nude mice, the BLI signal of PD2R cells dropped sharply to the lowest point after one day, and then stabilized at that level for an extended period (Supplementary Fig. S4A). Most of the mice injected with PD2R never developed overt bone metastasis (Supplementary Figs. S4B-D). Consistently, the Ki-67 staining of disseminated PD2R cells in the bone marrow indicated they were not proliferating. On the contrary, more than 30% of SCP28 cells were Ki-67 positive in the bone (Supplementary Figs. S4E&F).

We then overexpressed LTβ in PD2R cell line. These cells were IC injected into female athymic nude mice for bone metastasis progression. While the vector control cells did not form overt bone metastasis, most mice injected with LTβ OE cells developed macrometastasis (Figs. 4A&B). We further tested the role of LTβ in promoting PD2R bone metastasis progression in NSG mice, which are the most immunodeficient mice without T cells, functional macrophages, or natural killer (NK) cells (Shultz et al., 2007). PD2R-Vector cells developed moderate bone metastasis in NSG mice due to less immunosurveillance, OE of LTβ still significantly accelerated bone metastasis progression (Supplementary Fig. S4G). Severe bone degradation was also observed in the PD2R-LTβ group (Supplementary Fig. S4H and Supplementary Videos 4-5). Histological staining confirmed an increased bone metastatic area and an increased number of TRAP^+^ osteoclast cells in mice with LTβ OE (Supplementary Figs. S4H-K). These results suggest that LTβ strongly induced bone metastasis progression in indolent PD2R cells.

**Figure 4.**
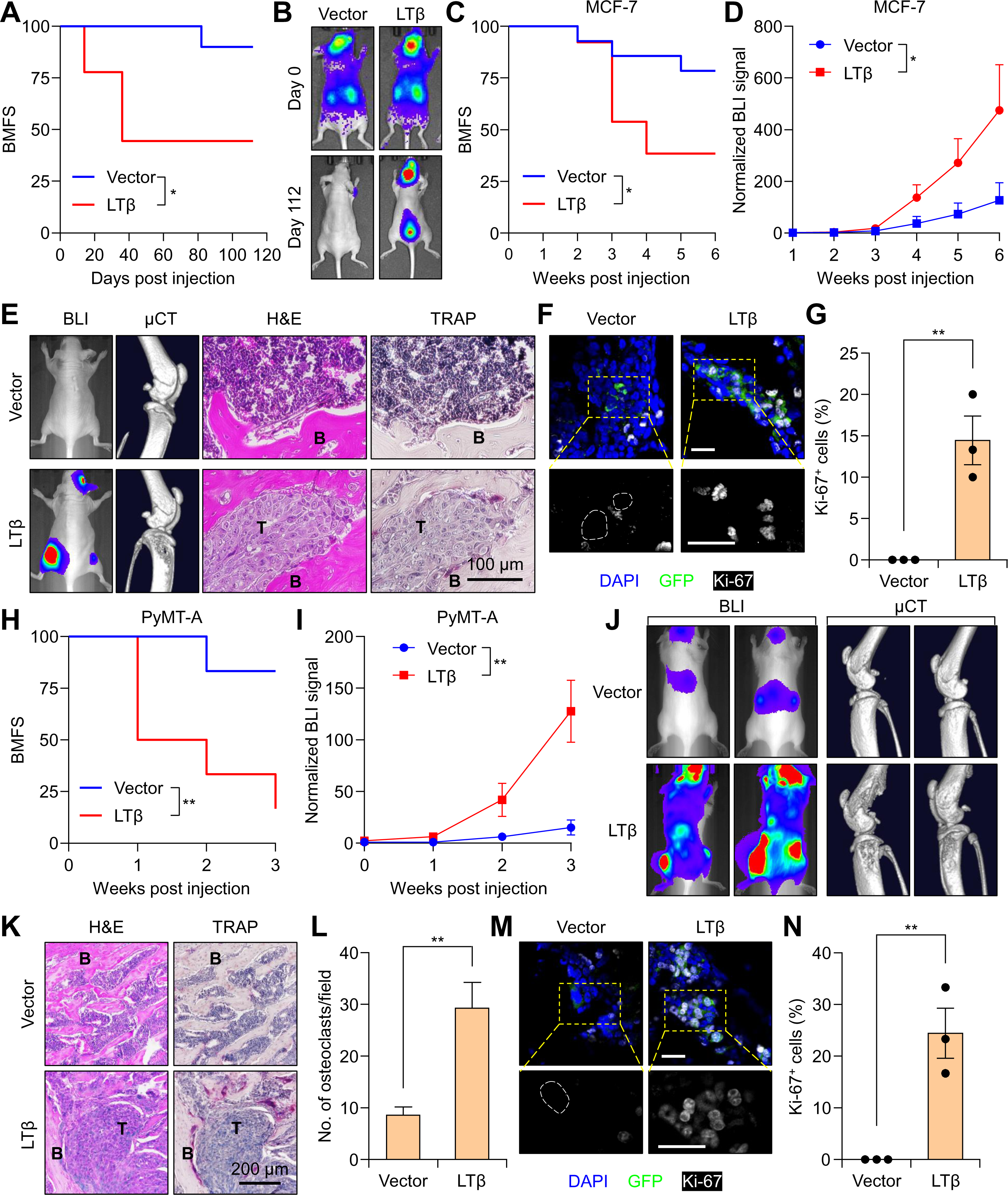
LTβ Reactivates Dormant Cancer Cells to Form Osteolytic Bone Metastasis. (**A**) PD2R cells with either Vector control or LTβ overexpression were utilized for bone metastasis assay. 10^5^ tumor cells were IC injected into 6 weeks old female nude mice. Bone metastasis burden was determined weekly by BLI imaging. Kaplan-Meier curve of mice bearing bone metastasis was presented. n = 10 mice for Vector control group, and n = 9 mice for LTβ group. *p < 0.05 by log-rank test. (**B**) Representative BLI images of mice bearing bone metastasis from experiment in **A**. (**C**) MCF7 cells with either Vector control or LTβ overexpression were utilized for bone metastasis assay. 10^5^ tumor cells were IC injected into 6 weeks old female nude mice. Bone metastasis burden was determined weekly by BLI imaging. Kaplan-Meier curve of mice bearing bone metastasis was presented. n = 14 mice for Vector control group, and n = 13 mice for LTβ group. *p < 0.05 by log-rank test. **(D)** Quantification of BLI signal from experiment in **C**. n = 14 mice for Vector control group, and n = 13 mice for LTβ group. Data presented as mean ± SD. *p < 0.01 by two-way repeated measures ANOVA. (**E**) Representative BLI, μCT, H&E staining, and TRAP staining images of bone metastasis bearing mice from experiment in **C**. B, bone tissue area; T, tumor area. Scale bar = 100 μm. (**F**) Representative IF images of Ki-67 in MCF7-GFP cells in bones. Similar experiment was performed as in **C**, a group of mice were sacrificed at Week 2 post injection for hind limb bone collection and IF staining against Ki-67. GFP: cancer cells. Scale bar = 20 μm, white dotted lines: nuclear border of GFP^+^ tumor cells. (**G**) Percentage of Ki-67^+^ cells detected by IF staining from experiments in **F**. n = 3 mice per each group. Data presented as mean ± SEM. **p < 0.01 by unpaired t-test. (**H**) PyMT-A cells with either Vector control or LTβ overexpression were utilized for bone metastasis assay. 10^5^ tumor cells were IC injected into 6 weeks old female FVB mice. Bone metastasis burden was determined weekly by BLI imaging. Kaplan-Meier curve of mice bearing bone metastasis was presented. n = 6 mice per group. **p < 0.01 by log-rank test. (**I**) Quantification of BLI signal from experiment in **H**. n = 6 mice per group. **p < 0.01 by two-way repeated measures ANOVA. (**J**) Representative BLI and μCT images of bone metastasis bearing mice from experiment in **H**. (**K**) Representative H&E staining and TRAP staining images of bone metastasis bearing mice from experiment in **H**. B, bone tissue area; T, tumor area. Scale bar = 200 μm. (**L**) Quantification of TRAP^+^ osteoclasts from experiment in **H**. n = 3 per group. Data presented as mean ± SEM. **p < 0.01 by unpaired t-test. (**M**) Representative IF images of Ki-67 in PyMT-A-GFP cells in bones. Similar experiment was performed as in **H**, a group of mice were sacrificed at Week 1 post injection for hind limb bone collection and IF staining against Ki-67. GFP: cancer cells. Scale bar = 20 μm, white dotted lines: nuclear border of GFP^+^ cells. (**N**) Percentage of Ki-67^+^ cells detected by IF staining from experiment in **M**. n = 3 mice for each group. Data presented as mean ± SEM. **p < 0.01 by unpaired t-test. **See also Supplementary Figure S4.**

To confirm the role of LTβ in bone dormancy reactivation, we utilized two additional dormant bone metastasis cell lines. MCF7, a human breast cancer cell line that is estrogen positive (ER^+^), is known to have little bone metastatic potential (Johnson *et al*., 2016). OE of LTβ in MCF7 cells significantly reduced the BMFS time and increased the bone metastasis burden compared to the vector control group (Figs. 4C&D). LTβ expression also significantly enhanced bone degradation, increased bone metastatic tumor areas, and increased the number of TRAP^+^ osteoclasts (Fig. 4E). IF staining of Ki-67 cell proliferation marker suggests that MCF7-Vector cells were quiescent in the bone, while LTβ-OE induced the cell cycle progression (Figs. 4F&G). PyMT-A is derived from *MMTV-PyMT* spontaneous mouse mammary tumors (Wan et al., 2014). Similar study as well as our preliminary test revealed that this type of cell line is dormant in the bone (Nobre *et al*., 2021). When cells were IC injected into FVB/N syngeneic mice, PyMT-A vector cells barely generated bone metastases. In contrast, OE of LTβ led to over 80% of mice developing strong bone metastasis (Figs. 4H&I). Consistent with the increased incidence of bone metastasis, OE of LTβ caused drastic bone degradation and increased the number of mature osteoclasts (Figs. 4J-L). IF staining of Ki-67 again suggested that PyMT-A-Vector cells were cell cycle arrested in the bone, while LTβ-OE induced the cell proliferation (Figs. 4M&N). In aggregate, our study utilized three distinct cell lines representing human triple-negative breast cancer, human ER^+^-breast cancer, and mouse mammary cancer cells and demonstrated that *LTB* is a dormancy reactivation gene.

### LTβ stimulates the expression of CCL2 and CCL5 in osteoblasts through NF-κB2 pathway

*LTB* expression was specifically induced only when cancer cells were in the bone microenvironment. This was evidenced by minimal expression when cells were collected from CL or FP, but high expression when cells were isolated from bone metastatic samples (4T1-D4, D10, D16, and 4T1.2-D4) (Fig. 5A). This result indicates that a bone microenvironment-specific signal may dictate *LTB* expression. Notably, *LTB* was already induced in the early seeding phase (D4), during which cancer cells mainly interact with osteoblasts (Wang *et al*., 2015; Zheng *et al*., 2017). Cancer cells were in close contact with osteoblasts at this stage (Fig. 5B). To explore this further, we examined *Ltb* expression when tumor cells (GFP-labeled) were co-cultured with either MC3T3 cells (mCherry-labeled), a mouse osteoblast cell line, or mouse mesenchymal stem cells (MSC, mCherry-labeled), progenitor cells for osteoblasts (Ren et al., 2008). In both cases, *Ltb* expression was dramatically induced in FACS sorted GFP-positive tumor cells (Fig. 5C), indicating that factors derived from osteoblastic lineage cells may be responsible for *LTB* expression in cancer cells.

**Figure 5.**
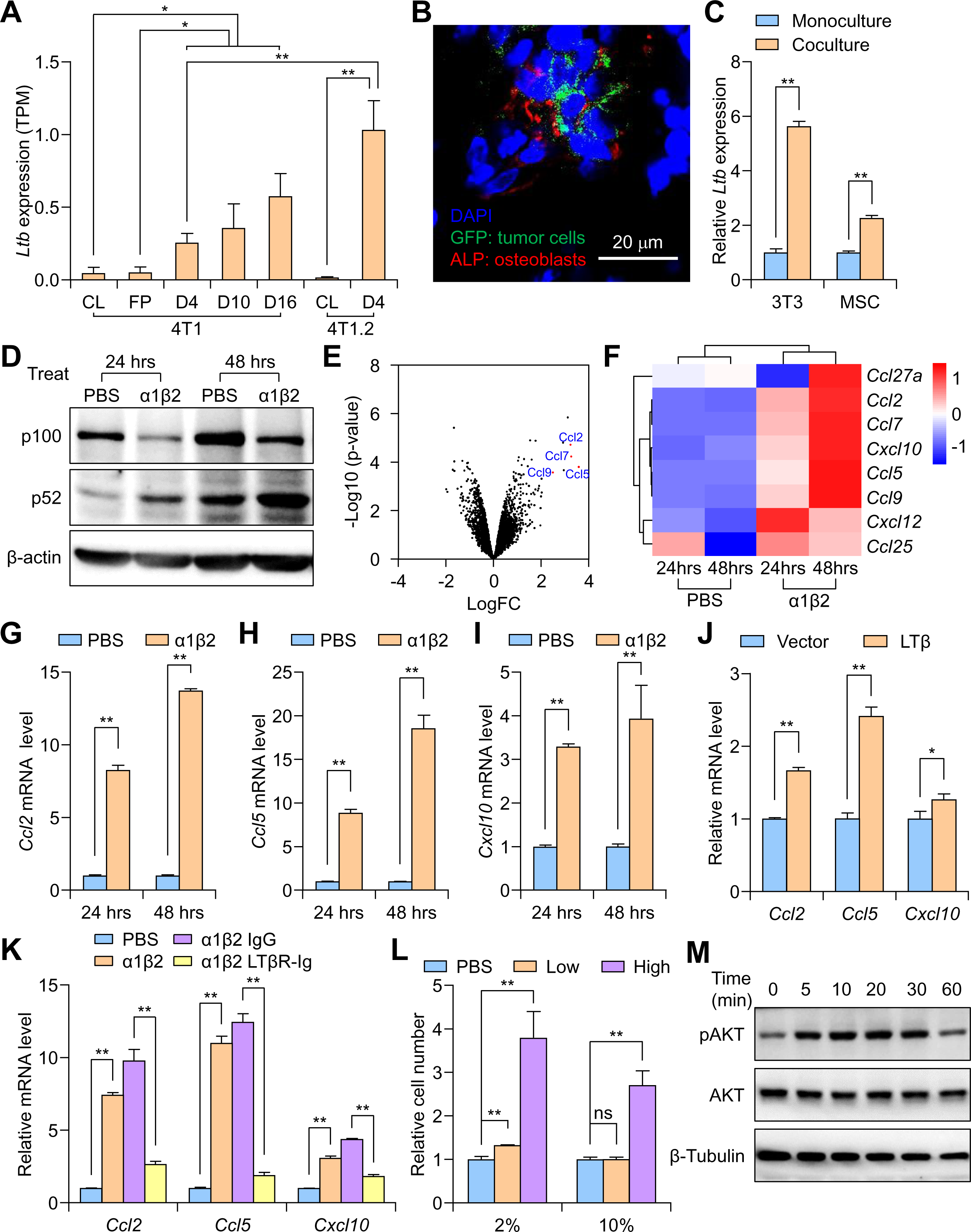
Tumor-derived LTβ Stimulates Chemokine Production in Osteoblasts through NF-κB2 Signaling. (**A**) The mRNA expression levels of *Ltb* in 4T1 cells from *in vitro* culture (4T1-CL), mammary fat pad injection (4T1-FP), and from different stages of bone metastasis (4T1-D4, D10, and D16) were quantified based on scRNA-seq results in Fig. 1B. The mRNA expression levels of *Ltb* in 4T1.2 cells from *in vitro* culture (4T1.2-CL) and early-stage (4T1.2-D4) bone metastasis were also analyzed. Data presented as mean ± SEM. *p < 0.05, **p < 0.01 by unpaired t-test. (**B**) GFP-labeled SCP28 were IC injected into nude mice. Four days post injection, mice were sacrificed to collect hindlimbs for IF staining. osteoblast cells were stained with anti-ALP antibody. Nuclei were counter-stained with DAPI (blue). Scale bar = 50 μm. (**C**) The mRNA expression level of *Ltb* in 4T1 cells as determined by q-PCR after co-culturing with MC3T3 or MSC. Data presented as mean ± SD. n =3. **p < 0.01. Significance was determined by unpaired t-test. (**D**) Representative immunoblot images of p100 and p52 in MC3T3 cells treated with LTα1β2 for 24 or 48 hrs. β-actin protein was used as an internal loading control. (**E**) Volcano plot depicting differentially expressed genes in LTα1β2-treated MC3T3 compared to that of PBS-treated cells. Ccl family members are highlighted. (**F**) Heatmap represents differentially expressed mRNAs for selected chemokines in LTα1β2-treated MC3T3 compared to that of PBS-treated cells. Scale bar indicates Z-scores. (**G**) The mRNA expression levels of *Ccl2* in PBS-or LTα1β2-treated MC3T3 were determined by qPCR. n = 3 per group. Data presented as mean ± SD. **p < 0.01 by unpaired t-test. (**H**) The mRNA expression levels of *Ccl5* in PBS-or LTα1β2-treated MC3T3 were determined by qPCR. n = 3 per group. Data presented as mean ± SD. **p < 0.01 by unpaired t-test. (**I**) The mRNA expression levels of *Cxcl10* in PBS-or LTα1β2-treated MC3T3 were determined by qPCR. n = 3 per group. Data presented as mean ± SD. **p < 0.01 by unpaired t-test. (**J**) The mRNA expression levels of *Ccl2*, *Ccl5, and Cxcl10* in MC3T3 cells after co-culturing with 4T1 cells were determined by qPCR. n = 3 per group. Data presented as mean ± SD. **p < 0.01 by unpaired t-test (**K**) MC3T3 cells were treated with PBS or LTα1β2. Additional decoy receptor, recombinant LTβR-Ig was further added into the LTα1β2-treated group, IgG was used as negative control. The mRNA expression levels of *Ccl2*, *Ccl5,* and *Cxcl10* in MC3T3 cells after these treatments were determined by qPCR. n = 3 per group. Data presented as mean ± SD. **p < 0.01 by unpaired t-test. (**L**) Relative tumor cell numbers were quantified by cell counting when SCP28 cells were cultured in the media containing 2% or 10% FBS and treated with 100 ng/ml or 200 ng/ml recombinant human CCL2 and CCL5 proteins. n = 3 per group. Data presented as mean ± SD. ns: p > 0.05, **p < 0.01 by unpaired t-test. (**M**) Representative immunoblot images of pAKT and total AKT in SCP28 cells treated with CCL2 and CCL5 proteins. β-Tubulin was used as an internal loading control. **See also Supplementary Figure S5.**

LTβ is a type II membrane protein that belongs to the TNF family and functions as a heterodimer with its α-subunit (Lu and Browning, 2014; Upadhyay and Fu, 2013). The classical receptor of LTβ is lymphotoxin-β receptor (LTβR), which is encoded by the *LTBR* gene (Ware, 2005). To investigate the mechanism by which LTβ mediates tumor-stromal interactions to promote bone colonization, we treated MC3T3 osteoblasts with recombinant human lymphotoxin α1/β2 protein (rhLTα1β2) and performed RNA-seq analysis. Heatmap and principal component analysis (PCA) showed that replicates of MC3T3 cells treated with rhLTα1β2 exhibited very similar gene expression patterns compared to the control group (Supplementary Figs. S5A&B). GO analysis revealed that rhLTα1β2 treated cells were enriched in immune response pathways, especially multiple pathways related to NF-κB signaling (Supplementary Fig. S5C). This finding is consistent with the known activation of NF-κB signaling by LTβ (Madge et al., 2008). Immunoblotting confirmed that the non-canonical NF-κB p52 protein level was significantly increased after rhLTα1β2 stimulation, further confirming activation of the NF-κB2 pathway (Fig. 5D).

Comparing the gene expression profiles between rhLTα1β2 treated cells *v.s.* control cells, we found that chemokine coding genes, including *Ccl2*, *Ccl5*, *Ccl7*, *Ccl9*, *Cxcl10*, and *Cxcl12*, were the most significantly changed ones (Figs. 5E&F and Supplementary Fig. S5D). We selected the Ccl2, Ccl5, and Cxcl10 genes for further study, as they were most upregulated and expressed at relatively abundant levels (Figs. 5G-I). Co-culturing of MC3T3 cells with 4T1 tumor cells showed similar results to the rhLTα1β2 stimulation experiment, confirming the induction of these chemokines by LTβ (Fig. 5J). Furthermore, LTβR-Ig, a decoy receptor for LTβ, was able to block the induced expression of *Ccl2*, *Ccl5*, and *Cxcl10*, indicating that the expression of these cytokines was dependent on LTβ-induced signaling (Fig. 5K). The activation of the NF-κB2 pathway in osteoblasts was further validated through IF staining of NF-κB2 downstream effector molecule CCL2 (Supplementary Fig. S5E). The presence of recombinant CCL2 and CCL5 proteins also significantly enhanced tumor cell proliferation, possibly through activation of the AKT pathway (Figs. 5L&M).

### LTβ-mediated tumor-osteoblast interaction enhances bone colonization and osteoclastogenesis

To investigate the role of LTβ in promoting tumor cell adhesion and bone seeding through osteoblasts, we conducted an *in vitro* cell adhesion assay. mCherry-labeled MC3T3 cells were cultured to confluence, and GFP-labeled SCP28 cells were added to allow adhesion for a short time. LTβ OE in SCP28 cells significantly increased adhesion to MC3T3 cells compared to vector control cells. LTβ-mediated tumor-osteoblast adhesion was blocked by LTβR-Ig (Figs. 6A&B and Supplementary Fig. S6A), which was confirmed in PD2R cells (Supplementary Fig. S6B). To examine whether LTβ promotes bone colonization *in vivo*, GFP-labeled SCP28 cells, with or without LTβ OE, were IC injected into nude mice. Four days later, hindlimb dissection and IF staining for tumor cells and osteoblasts detection revealed that LTβ facilitated tumor cell colonization, as significantly more tumor cells were detected in close proximity to bone osteoblasts when SCP28 cells were overexpressing LTβ (Fig. 6C).

**Figure 6.**
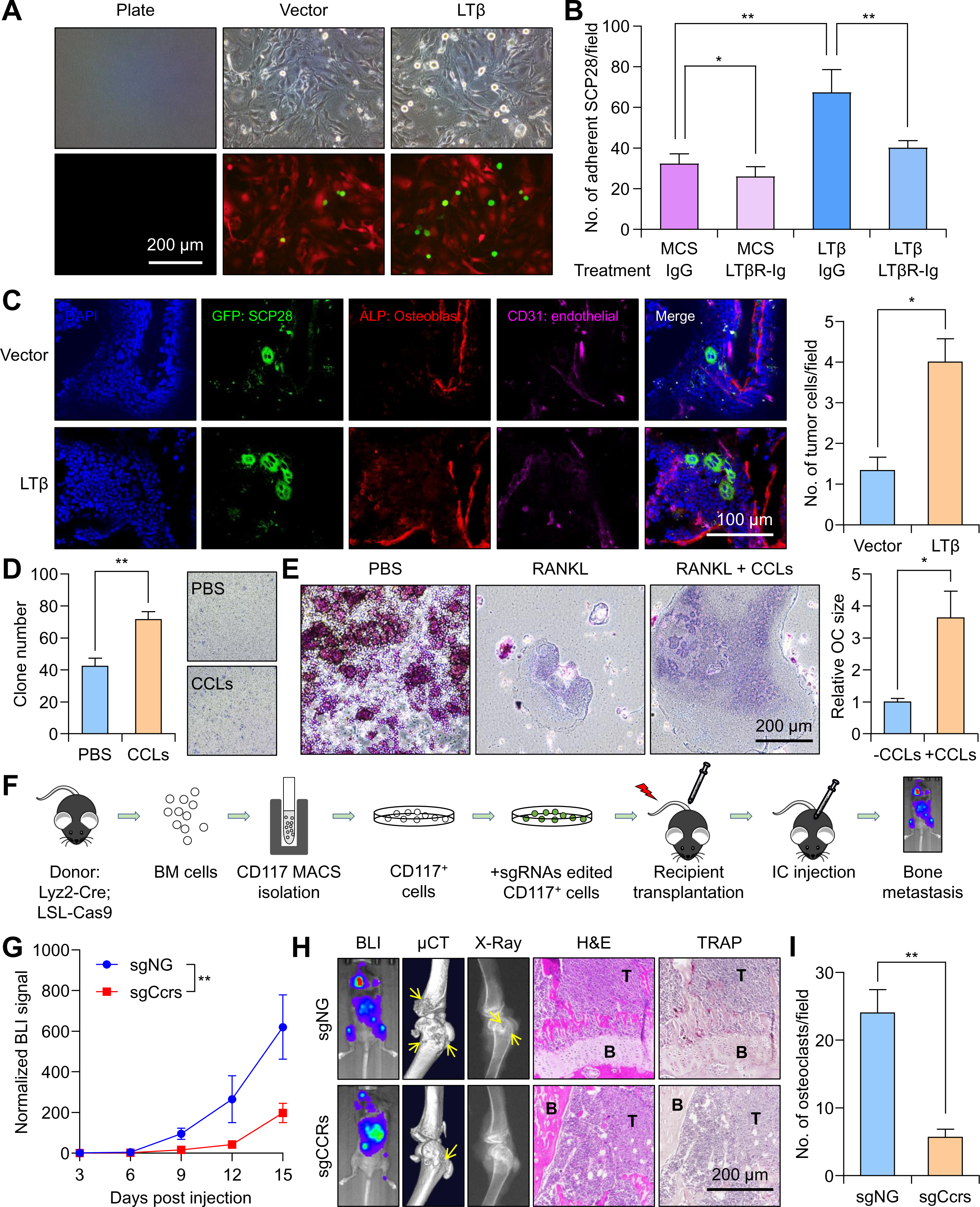
LTβ and CCL2/5 Promotes Bone Metastasis through Enhancing Cancer Cell-Osteoblast Adhesion and Osteoclast Differentiation. (**A**) mCherry-labelled MC3T3 cells were seeded onto cell culture plates to reach 100% confluency. GFP-labeled SCP28 cells with either-Vector or-LTβ expression were seeded on top of MC3T3 cells for 2 minutes. The number of adhered SCP28 cells was determined using direct microscopic imaging. Scale bar = 200 μm. (**B**) The experiment was performed similarly as in A, with additional groups treated with LTβR-Ig. The number of adhered SCP28 cells was quantified based on GFP signal. n = 6 per group. Data are presented as mean ± SD. *p < 0.05, **p < 0.01 by unpaired t-test. (**C**) SCP28 cells were IC injected into 6-week-old female nude mice. Mice were sacrificed 4 days post injection for hindlimb bone collection and IF staining. GFP: SCP28 cells; ALP: osteoblasts; CD31: endothelial cells. Scale bar = 100 μm. The number of GFP^+^ tumor cells per field was quantified in the right panel. n = 3 per group. Data are presented as mean ± SD. *p < 0.01 by an unpaired t-test. (**D**) Raw264.7 cells were used for transwell migration with CCLs (CCL2 and CCL5) added in the bottom of the transwell. Left panel: quantification of migrated cells. Right panel: representative images of migrated cells. n = 4 per group. Data are presented as mean ± SD. **p < 0.01 by unpaired t-test. (**E**) Raw264.7 preosteoclast cells were induced to differentiate to osteoclasts in the presence of RANKL. CCLs were added to one of the experimental groups. Osteoclasts were stained using an osteoclast staining kit. Scale bar = 200 μm. The relative size (diameter) of osteoclasts from RANKL group and RANKL + CCLs group were quantified in the right panel. n = 4 per group. Data are presented as mean ± SD. *p < 0.01 by unpaired t-test. (**F**) Schematic representation of the *in vivo* bone metastasis assay in CRISPR-Cas9-based, macrophage-specific knockout of CCR2/5 mouse model. Lyz2-Cre; LSL-Cas9 mice were sacrificed for bone marrow CD117^+^ cells collection (enriched for HSC cells). Bone marrow CD117^+^ cells were cultured *in vitro* and transduced with either the negative control sgRNAs or Ccr2/5 sgRNAs. The gene-edited CD117^+^ cells were transplanted into legally irradiated recipient mice to reconstitute their hematopoietic system. Mice were then used for *in vivo* bone metastasis analysis by IC injection of E0771-LTβ cells. (**G**) The bone metastasis burden from the experiment in **F** was monitored by BLI imaging and quantified. n = 3 mice for sgNG control group, and n = 4 mice for sgCcrs group. Data presented as mean ± SD. **p < 0.01, by two-way repeated measures ANOVA. (**H**) Representative BLI, μCT, X-ray, H&E staining, and TRAP staining images of bone metastasis from the experiment in **F**. B, bone tissue area; T, tumor area. Scale bar = 200 μm. (**I**) Quantification of TRAP^+^ osteoclasts from decalcified histological bone sections of hindlimbs from mice in **H**. n = 3 per group. Data presented as mean ± SEM. **p < 0.01 by unpaired t-test. **See also Supplementary Figure S6.**

Our results showed that LTβ promoted dormant cells to generate strong osteolytic bone metastases. Part of the reason might be that it facilitates tumor cell adhesion to the osteoblast niche. However, to generate osteolytic outgrowth, tumor cells need to recruit macrophages and enhance osteolytic bone degradation. Interestingly, CCL2 and CCL5, the main chemokines induced by LTβ in osteoblasts, have been reported to be involved in osteoclastogenesis (Binder et al., 2009; Lee et al., 2017). Indeed, CCLs (CCL2 and CCL5) induced macrophage recruitment in the transwell migration assay (Fig. 6D). Furthermore, supplementation with CCLs in an osteoclastogenesis assay also significantly enhanced osteoclast maturation (Fig. 6E). These results suggest that LTβ may be dependent on CCLs-mediated macrophage recruitment and osteoclastogenesis to promote bone metastasis. To test this possibility *in vivo*, we adopted a CRISPR-Cas9 delivery system to knock out CCR2 and CCR5 receptors specifically in macrophages and evaluated whether their depletion in macrophages blocked LTβ-induced bone metastasis (LaFleur et al., 2019; Wilkinson et al., 2019). As illustrated in Fig. 6F, Bone marrow CD117^+^ (BM CD117^+^) cells, which were enriched in hematopoietic stem cells (HSCs) from Lyz2-Cre (Cre driven by macrophage specific promoter); LSL-Cas9 mice were isolated and expanded *in vitro*. BM CD117^+^ cells were then transduced with lentivirus containing sgRNAs targeting CCR2 and CCR5. The knockout efficiency of these sgRNAs was first confirmed in mouse tumor cell lines by DNA sequencing and then directly in macrophages by FACS analysis (Supplementary Figs. S6C&D).

Furthermore, these BM CD117^+^ cells were differentiated into macrophages and used in *in vitro* osteoclastogenesis assay. CCR2 and CCR5 KO led to a dramatic decrease in the number of large, multi-nucleated osteoclasts (Supplementary Fig. S6E). BM CD117^+^ cells were then transplanted into lethally irradiated (5.5 Gy, two times) recipient mice to reconstitute their hematopoietic system. The re-constituted mouse hematopoietic system was determined by FACS analysis, and no obvious changes in the percentage of B, T, or myeloid cells were observed in re-constituted mice compared to wild type mice (Supplementary Figs. S6F&G). The recipient mice were rested for 5 weeks before IC injection of E0771 tumor cells (to match the B6 mouse background) to generate bone metastasis. In control mice that were transplanted with BM CD117^+^ cells of negative control sgRNA, E0771-LTβ led to strong osteolytic bone metastasis progression. In contrast, mice transplanted with BM CD117^+^ cells of CCR2 and CCR5 knockout generated much weaker bone metastasis (Fig. 6G). There was also a dramatic decrease in bone degradation and the number of TRAP^+^ osteoclasts (Figs. 6H&I). Thus, our results suggest that the LTβ-CCLs signaling axis enhances osteoclast differentiation to promote bone metastasis.

### LTβ is a potential therapeutic target for bone metastasis

To assess the therapeutic potential of targeting LTβ, we treated mice with recombinant LTβR-Ig decoy receptors to block LTβ-LTβR interactions and evaluated metastasis progression in multiple models. Mice were IC injected with SCP28-LTβ cells to generate bone metastasis. Three days later, LTβR-Ig treatment was initiated and continued until the experimental endpoint (Fig. 7A). BLI imaging showed that LTβR-Ig treatment significantly reduced the bone metastasis burden compared with that in the control group (Fig. 7B). LTβR-Ig treatment also prolonged BMFS while protecting the bone from degradation (Figs. 7C-E). Similar results were observed in PD2R-LTβ cells that were IC injected into NSG mice and treated with LTβR-Ig (Supplementary Figs. S7A-C). To rigorously evaluate the role of LTβ in bone metastasis, we turned to a humanized CD34^+^ (hu-CD34^+^) mouse model in which the mouse hematopoietic system was reconstituted by human HSC transfer. With this protocol, mouse blood circulation constituted a significant amount of human CD45^+^ cells (Supplementary Figs. S7D&E). When SCP28 cells were IC injected into these mice, LTβ OE significantly increased the number of bone-seeded cells within a week (Supplementary Figs. S7F&G).

**Figure 7.**
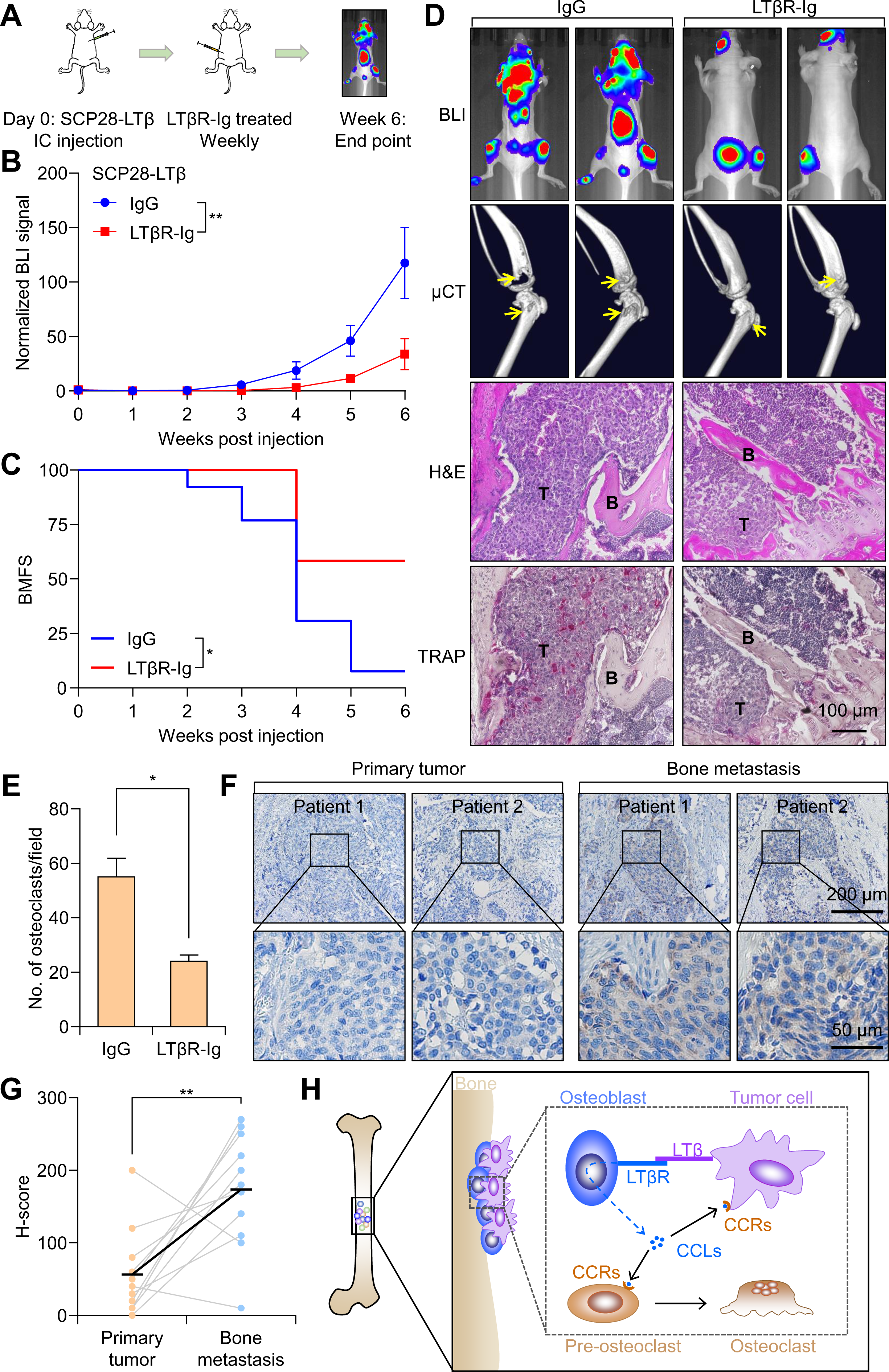
LTβ is a Therapeutic Target for Bone Metastasis. (**A**) SCP28-LTβ cells were IC injected into 6-week-old female nude mice. Mice were treated with LTβR-Ig recombinant protein three days later and continued until the experimental endpoint. (**B**) Bone metastasis burden was monitored by BLI imaging from experiment in **A**. n = 13 mice per group. Data are presented as mean ± SD. **p < 0.01 by two-way repeated measures ANOVA. (**C**) Kaplan-Meier BMFS curve of mice bearing bone metastasis in the experiment performed in **A**. n = 13 mice per group. *p < 0.05 by log-rank test. (**D**) Representative BLI, μCT, H&E staining, and TRAP staining images of bone metastasis from the experiment in **A**. B, bone tissue area; T, tumor area. Scale bar = 200 μm. (**E**) Quantification of TRAP^+^ osteoclasts from decalcified histological bone sections of hindlimbs from mice in **D**. n = 4 per group. Data presented as mean ± SEM. *p < 0.05 by unpaired t-test. (**F**) Representative IHC staining images of LTβ in paired human primary breast tumor and bone metastasis samples. Scale bar = 200 or 50 μm (magnified images). (**G**) IHC image scores (H-scores, see METHODS for detailed information) of LTβ in paired primary tumors and bone metastasis samples. Horizontal lines indicate the median values. n = 11 per group. **p < 0.01 by unpaired t-test. (**H**) Schematic model of LTβ in bone metastasis colonization and dormancy reactivation. When cancer cells arrive in the bone microenvironment, LTβ expression in cancer cells is induced by microenvironmental cues. LTβ binds to its receptor on osteoblasts, stimulates NF-κB2 activation, and induces chemokine (CCL2 and CCL5) production. CCL2 and CCL5 provide feedback to tumor cells to promote cell adhesion to the osteoblastic niche; they also recruit macrophages and enhance osteoclast differentiation. These effects facilitate tumor cell colonization of the bone and reactivate dormant tumor cells to generate osteolytic bone metastases. **See also Supplementary Figure S7.**

Our data strongly supports the role of LTβ in breast cancer bone metastasis. To confirm LTβ as a potential therapeutic target for patients with bone metastasis, we collected paired primary breast tumors and bone metastasis samples from 11 patients and performed Immunohistochemistry (IHC) staining against LTβ. The expression level of LTβ was consistently higher in bone metastasis samples than in paired primary tumor samples (Figs. 7F&G). These clinical data corroborate the prognostic value of *LTB*, as higher *LTB* expression is associated with shorter BMFS time in patients with breast cancer (refer to Fig. 2K), strongly suggesting that LTβ as a therapeutic target for patients with bone metastatic disease.

## DISCUSSION

Bone metastasis is a major challenge in breast cancer therapy, patients often develop it many years after considered clinical cure. This implies that bone-seeded cells remain quiescent in the bone microenvironment for an extended period before adapting to the niche and initiating overt metastatic growth. Understanding the signaling pathways involved in bone colonization is thus crucial for identifying therapeutic targets for the treatment of bone metastasis, and possibly dormancy and relapse. Although a few studies have begun to investigate tumor-stromal interactions critical for bone colonization, most rely on bulk sequencing results (Lu *et al*., 2011; Romero-Moreno et al., 2019). Consequently, the native signaling status of tumor cells during the early-seeding phase with single-cell resolution remains unknown. To address this, we used scRNA-seq analysis to explore the early bone-seeding signals. To our knowledge, our study generated the first set of scRNA-seq data for early-seeding bone metastatic tumor cells.

We conducted a comparative analysis of gene expression profiles between cells from early-stage bone metastasis (4T1-D4) and cells from *in vitro* culture (4T1-CL). Our analysis revealed that several GO terms related to secretion, cell adhesion, and cell-cell junctions were enriched, suggesting that the interaction between tumor and bone stromal cells plays a crucial role during colonization. To identify genes encoding membrane or secretion proteins that potentially mediate tumor-stromal interactions, we generated a mini cDNA library and functionally screened for genes that promote bone metastasis *in vivo*. To ensure statistical power, each cDNA was cloned with three independent barcodes, and only genes enriched from multiple mice were considered candidate genes. After clinical correlation analysis and additional functional validation, we identified LTβ as our top bone metastasis-promoting factor. LTβ is a member of the TNF superfamily and has not previously been reported to be involved in bone metastasis (Browning et al., 1993). The OE of LTβ increased osteolytic bone metastasis, while the KD of LTβ decreased the bone metastasis burden in multiple models. Notably, OE of LTβ in a comprehensive set of indolent bone metastatic cancer cells, including PD2R, MCF7, and PyMT-A cells, resulted in their reactivation and the generation of strong osteolytic bone metastasis. Thus, LTβ could possibly be considered as a strong dormancy reawakening factor.

In this study, we revealed the mechanism-of-action of LTβ in bone metastasis (Fig. 7H). Tumor-derived LTβ interacts with its receptor, LTβR, on osteoblasts, which enhances the tumor cell adhesion to bone osteoblasts. *In vivo* experiments showed that LTβ overexpression led to an increased number of tumor cells in close contact with ALP^+^ osteoblasts, confirming the importance of LTβ in mediating tumor-osteoblast interactions. LTβR activation on osteoblasts also stimulates the NF-κB2 pathway and resulting in the expression and secretion of chemokines including CCL2 and CCL5, which in turn recruit and promote the maturation of osteoclasts and facilitate bone degradation. Thus, LTβ helps bone metastasis progression in two important ways: mediating the initial seeding of tumor cells and promoting osteoclastogenesis to generate osteolytic outgrowth.

In addition to the induction of LTβ in the early-seeding phase of bone metastasis, multiple signaling pathways and interactions between tumor cells and stromal cells in the bone microenvironment may be involved in the progression of bone metastasis. Previous research has shown that the interaction between E-cadherin in tumor cells and N-cadherin in osteoblasts generates heterotypic cadherin junctions that facilitate bone metastatic colonization (Wang *et al*., 2015). Our single cell analysis of bone colonizing tumor cells suggest that successful colonization and metastatic outgrowth of tumor cells require the re-expression of epithelial marker genes, including E-cadherin. This suggests that LTβ and MET (E-cadherin expression) may synergistically promote tumor cell seeding in the bone. VCAM-1 has also been reported to promote the osteolytic expansion of indolent bone micrometastasis by interacting with α4β1-positive osteoclast progenitors (Lu *et al*., 2011). Our results confirm that LTβ can recruit macrophages and accelerate osteoclastogenesis, suggesting that LTβ and VCAM-1 might work together to generate an “osteolytic switch” that helps tumor cells to transit from interacting with osteoblasts to interacting with osteoclasts. Therefore, LTβ is both an “early-seeding molecule” and an “early-to-late switch molecule,” collaborating with other factors such as E-cadherin and VCAM-1 to facilitate tumor cell colonization and generate late-stage, osteolytic bone metastasis.

Based on our results, LTβ appears to be a promising therapeutic target for the treatment of bone metastasis. First, it has been shown to play a crucial role in promoting different stages of bone metastasis, including initial seeding, osteolytic bone metastasis, as well as possible dormancy reawakening. Second, our analysis of human bone metastasis samples showed that LTβ expression was significantly higher than that in paired primary tumors, providing further evidence of its role in bone metastasis. Third, we conducted proof-of-concept experiments using a LTβR-Ig decoy receptor to inhibit LTβ signaling and observed a significant reduction in bone metastasis progression across multiple models. Therefore, our findings support LTβ as a valuable therapeutic target for the treatment of bone metastasis, and further investigation in this direction is warranted.

## STAR METHODS

## RESOURCE AVAILABILITY

### Lead Contact

Further information and requests for resources and reagents should be directed to and will be fulfilled by the lead contact, Dr. Hanqiu Zheng.

### Materials Availability

The raw sequence data reported in this paper have been deposited in the Genome Sequence Archive in National Genomics Data Center, China National Center for Bioinformation / Beijing Institute of Genomics, Chinese Academy of Sciences (GSA: CRA007104) that are publicly accessible at https://ngdc.cncb.ac.cn/gsa.This paper does not report original code. Any additional information required to re-analyze the data reported in this paper is available from the lead contact upon request.

## EXPERIMENTAL MODEL AND SUBJECT DETAILS

### Animal Studies

All experimental protocols involving mice were approved by the Institutional Animal Care and Use Committee (IACUC) of Tsinghua University. All mice used in this study were female. BALB/c, FVB, and NU/NU mice were purchased from Vital-River. NSG mice were kindly provided by Dr. Xuerui Yang Lab in Tsinghua University. All mice were maintained under specific pathogen-free conditions at Tsinghua University. For orthotopic primary tumor formation, female BALB/c mice (4-6 weeks old) were anesthetized, and a small incision was made to reveal the mammary gland. A total of 10^5^ GFP-labeled 4T1 tumor cells suspended in 10 μL PBS were injected directly into the 4^th^ mammary fat pad. For intracardiac (IC) injection, tumor cells were harvested from sub-confluent cell cultures, washed with PBS, and suspended at 10^6^ cells/ml in PBS. Mice were anesthetized, 0.1 ml cells were injected into the left cardiac ventricle of 4-6 weeks old female mice using 26G needles. For MCF7 injection, self-made slow-release oestrogen (β-oestradiol 0.72 mg per implantation) pellets were implanted subcutaneously 3 days before cancer cell injection. Successful injection was confirmed by BLI imaging of evenly distributed bioluminescent signal in the entire body. For *in vivo* LTβR-Ig blockage assay, mice were IC injected with indicated cell lines in each experiment. Mice were then treated with 100 μg of LTβR-Ig or IgG protein per mouse three days post injection and continuously weekly until the experimental endpoint (Wu et al., 1999). The bone metastasis burden was monitored using BLI imaging, as mentioned in each experiment. At the experimental endpoint, the mice were euthanized, and bones were removed for μCT imaging and histological analysis.

### Cell Culture

HEK293T, 4T1, 4T1.2, PyMT-A, Raw264.7, E0771, SCP28, and PD2R cells were cultured in DMEM supplemented with 10% FBS and 1% penicillin-streptomycin. MCF7 cells were cultured in DMEM supplemented with 10% FBS, 1% penicillin-streptomycin, and 5 μg/ml insulin. MC3T3-E1 Clone #4 (denoted as MC3T3 for abbreviation in the text) was cultured in α-MEM with 10% FBS, 1% penicillin-streptomycin. Mouse mesenchymal stromal cell (MSC, isolated from BALB/c mouse) was cultured in DMEM supplemented with 10% FBS, 1% penicillin-streptomycin, and 20 ng/ml FGF. Cells were maintained at 37 °C in a humidified cell culture incubator containing 5% CO_2_ and 95% air. Cells were tested negative for mycoplasma contamination.

## METHOD DETAILS

### Virus Production and Generation of Stable Cell Lines

Lentiviral vectors were produced by co-transfecting HEK293T cells with lentiviral vectors and packaging plasmids psPAX2 and pMD2.G using Lipofectamine 2000, according to the manufacturer’s recommendations. Retroviral vectors were produced by co-transfecting HEK293T cells with retroviral vectors and packaging plasmids pCMV-Gag-Pol and pCMV-VSV-G using Lipofectamine 2000 transfection. Viral supernatants were collected at 48 hours post transfection, filtered through a 0.45 μm PVDF syringe filters (Cat# SLHV033RB, Millex). Viral transductions were performed for 12 hours in the presence of polybrene (8 μg/ml). Cells were expanded for another four days for FACS sorting or antibiotic selection at the appropriate concentrations.

### Generation of Knockdown and Overexpression Cells

Stable shRNA-mediated knockdown was achieved using the pLKO.1 lentivirus. The shRNA sequences used are listed in Supplementary Table S5. For shRNA knock down in the SCP28 cell line, shRNA viruses were generated and used to infect SCP28 cells as described above and subsequently selected with 2 μg/ml puromycin. For stable gene overexpression in the SCP28, MCF7, and PD2R human breast cancer cell lines, cDNA overexpression plasmid clones were picked from the barcoded mini-cDNA library (described below). Viruses were generated and used to transduce target cells, which were subsequently selected with 2 μg/ml puromycin. For stable gene expression in the PyMT-A and E0771 murine breast cancer cell lines, the full-length mouse *Ltb* gene coding sequence were cloned into the pMSCV-based retroviral plasmid. Viruses were generated and used to infect target cells as described above, and cells were selected with 2 μg/ml puromycin.

### Labeling of 4T1 and 4T1.2

We utilized pLVX-Ubc-Luciferase-GFP lentivirus and pMSCV-mCherry retrovirus to label the cells to facilitate in *vivo* BLI and cell recovery for scRNA-seq. pLVX-Ubc-Luciferase-GFP was a gift from Dr. Yonghui Zhang in Tsinghua University. To generate pMSCV-mCherry plasmid, mCherry fragment was PCR amplified using primers mCherry-XhoI-F and mCherry-SalI-R (listed in Supplementary Table S3), and inserted into pMSCV-Hygro at XhoI and SalI sites. 4T1 and 4T1.2 cells were then transduced with these two viruses and recovered for 4 days before FACS sorting to collect GFP^+^/mCherry^+^ double positive tumor cells.

### Cancer Cell Preparation for scRNA-seq

*In Vitro* Cultured Cells (4T1 and 4T1.2 cells used here were GFP^+^/mCherry^+^ cells.) 4T1 and 4T1.2 cells were cultured to about 80% to 90% confluency. Cell medium was removed and cells were washed twice with PBS. Cancer cells were trypsinized with 0.25% Trypsin-EDTA. Cancer cells were then washed with PBS and suspend in FACS buffer. Single cell suspension was filtrated through a 70 μm nylon cell strainers (Falcon, Cat# 352350) and FACS sorted by BD Influx flow cytometer to make sure only one cell was presented in each well of 96-well plate.

#### Mammary Fat Pad Injection-based Primary Tumor Cells

10^5^ 4T1 tumor cells were MFP injected into BALB/c mice. Mice were sacrificed at Day 22 and tumor tissues were dissected out for single cell solution preparation. Tumor cells were sorted by BD Influx flow cytometer to make sure only one cell was presented in each well of 96-well plate. The detailed protocol for the preparation of single cell suspension were performed similar as bone metastasis-based single cell capture described below.

#### Bone Metastasis Tumor cells

10^5^ 4T1 or 4T1.2 cells were IC injected into BALB/c mice. At Day 4, Day 10, and Day 16 post injection, a group of mice were sacrificed and hindlimb bones were dissected out. Hindlimb bones were shearing into small pieces and digested for 1 hour at 37 °C in DMEM medium containing 10% FBS, 300U/ml type I collagenase (Gibco, Cat# 17100017) and 0.1 mg/ml DNase I (Sigma, Cat# 260913). Digested tissues were filtrated through a 70 μm nylon cell strainers (Falcon, Cat# 352350). The cells suspensions were depleted of red blood cells using RBCs lysis buffer (Beyotime Biotechnology, Cat# C3702). Single cell suspension was filtrated through a 70 μm nylon cell strainers (Falcon, Cat# 352350) and FACS sorted by BD Influx flow cytometer to make sure only one cell was presented in each well of 96-well plate.

### Single-cell RNA-seq Library Preparation and Sequencing

Reverse transcription and cDNA amplification were carried out using a modified Smart-seq2 protocol (Picelli *et al*., 2014). Single cells were sorted by FACS as described above into 96-well plates containing 0.1 μl of RNase inhibitor (Clontech, Cat# 2313A), 1.9 μl of 0.2% Triton X-100 (Sigma, Cat# T9284), 1 μl of 10 µM oligo-dT primer (5′-AAGCAGTGGTATCA ACGCAGAGTAC T30VN-3′) and 1 μl of dNTP mix (10 mM each; Fermentas, Cat# R0192) in each well. Plates were either stored at −80 °C or processed immediately. Reverse transcription, PCR preamplification, PCR purification were performed following Picelli’s protocol (Picelli *et al*., 2014). As mRNA quality control, 1 ul of each PCR product was used as template for qPCR detection of mCherry expression and samples with Ct value higher than 30 will be discard. PCR products were purified with 1 × Agencourt Ampure XP beads (Beckman Coulter, Cat# A63881) and the final product was reconstituted in 20 µl TE buffer. The concentration of cDNA was measured by Qubit™ 1× dsDNA High Sensitivity (HS) Assay Kit (Invitrogen, Cat# Q33230). Tagmentation and NGS library was carried out by using TruePrep DNA Library Prep Kit V2 for Illumina (Vazyme, Cat# TD503) following the manufacturer’s instructions. The purified library was qualify controlled by measuring the fragment size distribution with an Agilent HS DNA BioAnalyzer Chip and the concentration of library was measured with Qubit™ 1× dsDNA High Sensitivity (HS) Assay Kits following the manufacturer’s instructions. Single cell cDNA libraries were sequenced on an Illumina HiSeq X Ten platform to obtain paired-end 150-bp reads.

### Processing of scRNA-seq Data

#### Generating Counts

Raw reads were first analyzed for quality control using FastQC (v0.11.7) (Andrews, 2010). Adaptor sequences of raw reads were removed by using TrimGalore (v0.6.6) (https://github.com/FelixKrueger/TrimGalore). Trimmed reads were mapped to a concatenated mouse genome (mm10) with STAR (v2.6.0c) with default parameters to result SAM format files (Dobin et al., 2013). Aligned reads stored in SAM format were transformed to generate BAM files using SAMtools (v1.7) (Danecek et al., 2021). The gene-level read count matrix was summarized from BAM files using FeatureCounts program (v1.6.2) (Liao et al., 2014). Subsequent data analysis was carried out in R 3.6.2 and the Seurat package (v3.1.2) (Stuart et al., 2019).

#### Quality Control and Normalization

The gene counts per cell, the percentage of mitochondrial transcripts, the percentage of ERCC RNA spike-in, the percentage of α and β-globin genes were computed using the functions of the Seurat package. Quality controls were performed following: a. Cells with luciferase or GFP sequence detected were kept for further analysis. b. Cells with transcripts counts lower than 5 × 10^5^ or higher than 1 ×10^7^ were discarded. c. Single cell libraries with number of detected gene lower than 5,000, mapping rate lower than 40% or the percentage of mitochondrial transcripts higher than 10% were discarded before further analysis. d. Libraries with the percentage of α and β-globin genes higher than 1% were also discarded. 424 single cells passed quality controls, and were used for further analysis. The gene expression level was normalized as log_2_(TPM/10 + 1), in which TPM is the abbreviation for transcripts per million, and adding one to avoid negative expression values.

#### Identification of Variable Genes and Dimensionality Reduction

Highly variable genes were identified with the function “FindVariableGenes” in Seurat following default parameters. These variable genes were then selected for downstream dimensionality reduction and clustering. Principle component analysis was performed on this scaled dataset. Cells were clustered based on the PCA scores of the first 20 principal components. The clustering of the data was performed based on the Seurat function “FindClusters” with a resolution of 1.6 and cells were clustered into 10 groups. In order to visualize the data, a “RunTSNE” parameter was used. The corresponding relationship between cluster and cell type is listed in Supplementary Table S1.

#### Identification of Marker Genes

The marker genes for interested population were identified using the FindMarkers function in Seurat. For example, to identify highly expressed genes in 4T1-D4 than 4T1-CL, we used FindMarkers, ident.1 = c (1, 4), ident.2 = 3, only.pos = TRUE, min.pct = 0.25 parameters. Where the value of ident here is corresponding with cluster number shown in Supplementary Table S1. Similarly, for 4T1.2-D4 *v.s.* 4T1-D4, we used 8 *v.s.* (1,4); for 4T1-CL *v.s.* other cells from 4T1, we used 3 *v.s.* (0, 1, 4, 5, 6, 7, 9); for 4T1-FP *v.s.* other cells from 4T1, we used 7 *v.s.* (0, 1, 3, 4, 5, 6, 9); for 4T1-BM *v.s.* other cells from 4T1, we used (0, 1, 4, 5, 6) *v.s.* (3, 7). As shown in Table S1, cells from 4T1-D4 were divided into cluster 1 and cluster 4. Marker genes were identified by using 1 or 4 as ident.1 or ident.2 in parameters, respectively.

#### Gene Ontology (GO) Enrichment Analysis

Previous identified sets of marker genes were used to perform gene ontology enrichment analysis using enrichGO function of clusterProfiler (v3.14.3) (Yu et al., 2012) in R with parameters of pAdjustMethod = “BH”, pvalueCutoff = 0.05, qvalueCutoff = 0.2, keyType = “ENTREZID”. Selected enriched GO terms were then plotted by using GraphPad Prism 8.

#### Single-cell Trajectories in the Seudotime Analysis

4T1 cells (excluding 4T1.2 data set) were selected and t-SNE was performed again as previously described. Gene expression matrix was used to study pseudotime trajectories of cells by using monocle2 (v2.14.0) (Trapnell et al., 2014) in R.

#### EMT Analysis Utilizing scRNA-seq Data

Expression matrix of single cells from FP, CL and different stages of bone metastasis was extracted from Seurat in log_2_(TPM/10 + 1) format. The expression panel of epithelial marker genes (*Epcam*, *Cdh1*, *Cldn3*, *Cldn4*, *Krt19*, and *Krt7*) and mesenchymal marker genes (*S100a4*, *Fn1*, *Vim*, and *Snai1*) in single cells were selected for heatmap construction by utilizing pheatmap (v1.0.12) package in R. Epithelial score was calculated by the averaged expression level of *Epcam* and *Cdh1*.

### cDNA Library Construction

#### Candidate Gene Selection and Mini-cDNA Library Construction

pLVX-CMV-MCS-mPGK-puro was utilized in as the lentiviral vector for cDNA cloning. To facilitate the generation of barcoded cDNA library, random 10N sequence were inserted into the vector at XhoI and AgeI restriction sites. Transformed *E.coli* bacteria lawn were picked for plasmid extraction. This pooled plasmid backbone was termed as pLVX-CMV-MCS-10N-mPGK-puro, and was utilized to generate random barcoded cDNAs for library construction.

Top 200 highly expressed genes in cells by comparing 4T1-D4 *v.s.* 4T1-CL and top 200 highly expressed genes by comparing 4T1.2-D4 *v.s.* 4T1-D4 were selected for analysis. The sub-cellular location of candidates was examined at Uniprot website (Apweiler et al., 2004). Membrane and secreted proteins were selected for cDNA cloning. The corresponding candidate human cDNAs were PCR amplified by using the following primer (Forward, CTAGAGGATCTATTTCCGGTGAATTCGCCACC-ATG-first 21-necleotides of CDS, Reverse, TCGTCGTCATCCTTGTAATCGGCGGCCGC-stop codon and last 21-necleotides of CDS). Amplicons were purified by agarose gel extraction and inserted into pLVX-CMV-MCS-10N-mPGK-puro at EcoRI and NotI sites by using NEBuilder HiFi DNA Assembly Master Mix (NEB Cat# E2621L). At least 3 individual *E.coli* clones were picked for plasmid extraction and plasmids were sequenced with CMV-Forward and mPGK-Reverse primers (See Key resource table) to confirm the cDNA sequence and the uniqueness of each barcode. 92 genes were chosen for cDNA library construction and 84 of them were cloned successfully. Each cDNA has three independent barcodes were selected to enhance the robustness of the *in vivo* screening. Totally, 254 independent cDNA vectors with one empty vector control were contained in this mini-cDNA library. Concentration of each plasmid was measured by nanodrop and plasmids were pooled into the library following the referred literature (Cante-Barrett et al., 2016).

#### *In Vivo* cDNA Library Screening and Candidate Gene Selection

Lentiviral cDNA library was packaged following methods described above and SCP28 cells were infected at MOI of 0.3 to ensure a single cDNA integration per cell. Cells were then selected with puromycin and recovered for a couple of days. Stably transduced cells were then IC injected into 4-6 weeks old female nude mice (3 × 10^5^ cells per mouse for 10 mice) for bone metastasis. The coverage rate for each bar code was more than 10,000 times. Six weeks post injection, mice were dissected, hindlimb bones were dissected out and cut into small pieces. Genomic DNA was purified by Phenol/chloroform extraction (Green and Sambrook, 2017). Genomic DNA was used as template for two rounds of PCR prior to next generation sequencing. In the first round of PCR, we used primer set of NGS-F and NGS-R (sequences listed in Supplementary Table S4) to amplify the barcode sequences. After PCR amplification, DNA fragments were purified as template for second round PCR to add index for NGS using primer listed in Supplementary Table S4. Samples were sequenced on a paired-end 150-bp run in Illumina HiSeq X Ten platform. Barcode sequences were extracted from Illumina sequence reads after removing common sequences around the barcodes. Absolute number of each barcode and total barcodes were counted, barcode and gene enrichment analysis were performed using software drugZ (v1.1.0.2) (Colic *et al*., 2019).

### RNA Extraction and RNA-seq Analysis

Total RNA was isolated from cells using RNAiso Plus Regent (Takara, Cat# 9108) following the manufacturer’s instructions. The NEBNext Ultra II Directional RNA Library Prep Kit for Illumina (NEB, Cat# E7765) was used to generate sequencing libraires from 1 μg of total RNA. All library preparations were conducted according to the manufacturer’s instructions. Paired-end 150bp sequencing was performed using an Illumina NovaSeq platform. Quality control, sequence mapping, and counts for each gene were performed following similar protocols as in scRNA-Seq. Gene expression matrix was normalized by using EdgeR (v3.30.0) package (Robinson et al., 2010) and carried out as counts per million (CPM) form.

### Quantitative PCR

Total RNA was isolated from cells using RNAiso Plus Regent (Takara, Cat# 9108) following the manufacturer’s instructions. RNA was reverse transcribed into cDNA by using ReverTra Ace^®^ qPCR RT Master Mix, Oligo (dT) (TOYOBO, Cat# FSQ-201). qPCR was performed on ABI QuantStudio 3 series PCR machine (Applied Biosystem) using the Power Green qPCR Mix (DONGSHENG BIOTECH, Cat# P2105). qPCR primers were listed in Supplementary Table S5. The gene-specific primer sets were used at a final concentration of 0.4 μM. Relative expression values of each target gene were normalized to *GAPDH* mRNA level. See Supplementary Table S6 for oligo sequences of each gene.

### Western Blot Analysis

Lysis buffer (50mM Tris-HCl pH 7.4, 150mM NaCl, 1m MEDTA, and 1% NP-40) was used to lyse the cells. Samples were heat denatured and equally loaded, separated on a 10% SDS-PAGE gel, transferred onto a PVDF membrane (Millipore), and blocked with 5% skim milk. Primary antibodies for immunoblotting included: α-β-actin (1:10,000 dilution, Abcam, Cat# ab6276, clone AC-15), α-NFκB p52 (1:1,000 dilution, Santa Cruz Biotechnology, Cat# sc-7386, clone C-5), α-Phospho AKT (1:1,000 dilution, Cell Signaling Technology, Cat# 4060), α-AKT (1:1,000 dilution, Cell Signaling Technology, Cat# 4691). Secondary antibodies used in experiments were included as following: Goat α-Mouse IgG (H&L)-HRP conjugated (1:10,000 dilution, Easybio, Cat# BE0102-100), Goat α-Rabbit IgG (H&L)-HRP conjugated (1:10,000 dilution, Easybio, Cat# BE0101-100). The chemiluminescence signals were detected by ECL substrate (Tanon, Cat# 180–5001) on ChampChemi Digital Image Acquisition Machine (Beijing Sage Creation Science Co, LTD).

### Tumor Cell Adhesion Assay

MC3T3 cells were cultured in 12-well plates until 100% confluency and pre-treated with serum-free DMEM containing 1 μg/ml LTβR-Ig protein or control IgG 15 minutes before adding tumor cells. GFP-labeled SCP28 or PD2R tumor cells overexpressing LTβ were non-enzyme digested off from 10-cm plate and washed with PBS. Tumor cells were diluted to 1 × 10^6^ cells/ml in DMEM medium without FBS. 5 × 10^5^ tumor cells were added into wells with pre-cultured MC3T3 cells, and incubated for 2 minutes. After incubation, the cell medium and unattached tumor cells were washed off. The remaining adherent cells were utilized for microscopy imaging and direct cell counting.

### Luciferase Assay

The cells were cultured in 24-well plates. At the end of the cell culture period, 100 μL of luciferase lysis buffer (2 mM EDTA, 20 mM DTT, 10% glycerol, 1% Triton X-100, 25mM Tris-base, pH 7.8) was added to each well. The cells were incubated in lysis buffer at room temperature for one hour with rotation at approximately 100 rpm. 25 μL cell lysate was transferred into a 96-well white plate, and 75 μL luciferase assay buffer (25 mM glycylglycine, 15 mM potassium phosphate, 4 mM EGTA, 2 mM ATP, 10 mM DTT, 1 mM D-luciferin, 15 mM MgSO_4_, pH 7.8) was added for immediate quantification using an EnVision^®^ Multimode Plate Reader (PerkinElmer).

### Mouse CD117^+^ Bone Marrow Cell Isolation, Gene Editing and Transplantation

CD117^+^ bone marrow cell (enriched for HSC cells) culture was performed according to a previously described method (Wilkinson et al., 2020). Briefly, mouse bone marrow cells were isolated from the tibia of Lyz2-Cre; LSL-Cas9 mice in C57/BL6 background. CD117^+^ cells were enriched using an autoMACS system (Miltenyi Biotec) with CD117 MicroBeads (Miltenyi, Cat# 130-091-224). These cells were cultured in fibronectin pre-coated 48-well plates with medium composed of F-12 medium (Gibco, Cat# 11765-054), 1x ITSX (Gibco, Cat# 51500-056), 10mM HEPES (Gibco, Cat# 15630-080), 1 mg/ml PVA (Sigma, Cat# P8136), 1% penicillin-streptomycin (Gibco, Cat# 15070-063), 1x GlutaMax (Gibco, Cat# 35050-061), 100 ng/ml mouse TPO (Novoprotein, Cat# AF-315-14), and 10 ng/ml mouse SCF (Novoprotein, Cat# AF-250-03). Cells were maintained at 37 °C in a humidified cell culture incubator containing 5% CO_2_ and 95% air. For gene knock out in CD117^+^ bone marrow cells, the lentivirus was packed as described above and concentrated using PEG-8000. GFP^+^ cells were sorted using FACS and cultured for bone marrow transfer. For CD117^+^ bone marrow cell transplantation, mice were irradiated with 5.5 Gy twice at 3 hours interval, and 1.5 x 10^6^ CD117^+^ cells were injected into each mouse by tail vein injection. Five weeks after transplantation, cancer cells were IC injected into mice.

### Humanized Mouse Model

Reconstitution of humanized mice was performed as previously described (Cao et al., 2021). Briefly, human CD34^+^ cells purified from cord blood were intravenously injected into female NSG mice (4 weeks old), which were irradiated 8 hours before transfer. Mice were maintained for approximately 12 weeks to allow for human immune reconstitution before cancer cell injection.

### Histomorphometric Analysis of the Bone Samples

Hindlimb bones were excised from mice at the end point of each experiment, immediately after the last *in vivo* BLI imaging. The tumor-bearing bones were fixed in 10% neutral-buffered formalin, decalcified in 10% EDTA for 2 weeks, and embedded in paraffin for H&E or TRAP staining (Kos et al., 2003). Histomorphometric analysis was performed on H&E stained images using a Nikon microscope and NIS-Elements BR software version 4.30. The osteoclast number was assessed as multi-nucleated TRAP^+^ cells and reported as the number of osteoclasts per field.

### X-Ray and μCT Imaging

Femurs and tibias were scanned using a Quantum GX microCT Imaging System (PerkinElmer) at the Laboratory Animal Resource Center, Tsinghua University. “Live Mode” was used and images in “Xcapture window” were saved as X-Ray images. μCT scanning was performed with the voltage of 90 kV, field of view (FOV) of 72 mm, and images were acquired at scan mode of high resolution for 4 minutes. The images were reconstructed to obtain a 3D structure for further analysis using Quantum GX microCT software (PerkinElmer).

### Immunofluorescence Staining of Bone Samples

For immunofluorescence staining of bone samples, hind limb bones were excised from the mice at the experimental endpoint and immediately fixed in 4% freshly prepared paraformaldehyde. Non-decalcified bone samples were frozen in embedding media and sectioned with Leica CM3050S Research Cryostat at 20 μm with Cryofilm type IIIC (Section-Lab, Japan). Sections were stained with the respective primary antibodies and fluorophore-labelled secondary antibodies. ALP antibody, 1:100, Cat# MAB29091 (R&D Systems). Ki-67 antibody, 1:100, Cat# ab15580 (Abcam). CCL2 antibody, 1:100, Cat# ab308522 (Abcam). Images were captured using an Olympus FV3000 confocal microscope at the Center of Biological Analysis, Tsinghua University.

### Immunohistochemistry Staining for Clinical Patient Samples

Paired primary breast cancer and bone metastasis samples were acquired from the Department of Orthopedic Oncology, the Affiliated Hospital of Qingdao University. All patient samples were de-identified for further transfer and analysis. Formalin-fixed, paraffin-embedded primary and bone metastasis sections were analyzed for LTβ expression. In brief, tissue sections were incubated in 0.01 M citrate buffer (pH 6.0) buffer at 95°C for 40 minutes to retrieve antigenicity. After treated with 3% H_2_O_2_ for 30 min to block endogenous peroxidase, the slides were incubated with α-LTβ antibody (Atlas Antibodies, HPA048884) at a 1:100 dilution for 1 hour at room temperature. After washing with PBS, the slides were then incubated with HRP-conjugated goat α-rabbit IgG antibody for 30 min at room temperature. Sections were stained with DAB and then counter-stained with hematoxylin according to manufacturer’s instructions. Images were obtained using a microscope (Nikon Eclipse Ti2). Intensity scoring was performed following a common four-point scale. 0 represents no staining, 1 represents low but detectable degree of staining, 2 represents clearly positive staining, and 3 represents strong expression. Expression was quantified as H-Score, by combining the results of staining intensity and the percentage of stained cells using the following standard formula: [1 × (% cells 1+) + 2 × (% cells 2+) + 3 × (% cells 3+)].

### Quantification and Statistical Analysis

Results were reported as mean ± SD (standard deviation) or mean ± SEM (standard error of the mean), as indicated in the figure legends. Unpaired t-test or two-way ANOVA analysis was used to calculate the statistical significance for the difference in a particular measurement between groups. p value less than 0.05 was considered to be statistically significant.

## ACKNOWLEDGMENTS

We thank J. Massague and Y. Kang for the generous gifting of multiple bone metastatic cell lines, including SCP28 and PD2R cells. We thank Y. Kang for the helpful discussions. We thank all members of the Zheng Laboratory for their helpful discussions and technical assistance. We thank the Technology Center for Protein Sciences at Tsinghua University for FACS support. We also thank the Laboratory Animal Research Center and the Center of Biomedical Analysis at Tsinghua University for providing the animal support. This study was partially supported by the National Key Research and Development Program of China (2020YFA0509400 to H.Z.), the National Science Foundation of China (81772981 and 81972462 to H.Z.), the Tsinghua University Initiative Scientific Research Program, and the Tsinghua-Peking Center for Life Sciences.

## AUTHOR CONTRIBUTIONS

X.W. and H.Z. designed and performed experiments. T. L. and J. W. provided technical support for the scRNA-seq experiments. T.Z. and X.W. performed the scRNA-seq analysis and provided in-house bioinformatics analysis. B.Z., Z.Z., and B.Y. collected and provided paired clinical breast cancer patient samples from primary breast cancer and bone metastatic tumor tissues. Y.L. and D.P. provided critical reagents and mouse models for mouse HSC isolation, knockout, and transplantation experiments. Y.L. and Y.-X.F. provided the humanized mouse model for this study. G.X., L.Z., Y.W., Q.S., H.Y., H.H., and X.L. provided experimental assistance for molecular cloning and animal works. H.Z. developed the concept, designed the experiments, and supervised the overall study. X.W. and H.Z. wrote and revised the manuscript.

## DECLARATION OF INTERESTS

The authors declare no conflict of interests in this study.

## Notes

### Competing Interest Statement

The authors have declared no competing interest.

