## Supplemental Figure 1 - 7 for "Lymphotoxin*-*β Promotes Bone Colonization and Osteolytic Outgrowth of Indolent Bone Metastatic cells of Breast Cancer"

**SUPPLEMENTARY INFORMATION**

**Supplementary figures**

Figure S1 is related to Figure 1.

Figure S2 is related to Figure 2.

Figure S3 is related to Figure 3.

Figure S4 is related to Figure 4.

Figure S5 is related to Figure 5.

Figure S6 is related to Figure 6.

Figure S7 is related to Figure 7.

**Supplementary tables:**

Table 1. The correlation between cell types and clusters in t-SNE plot.

Table 2. Candidates for cDNA library construction.

Table 3. DNA Oligos for plasmid construction.

Table 4. Oligos for NGS library construction.

Table 5. shRNA sequences.

Table 6. Oligos for qPCR.

**Supplementary videos:**

Supplemental Videos S1-3 related to Figure 3.

Supplemental Videos S4-5 related to Figure S4.

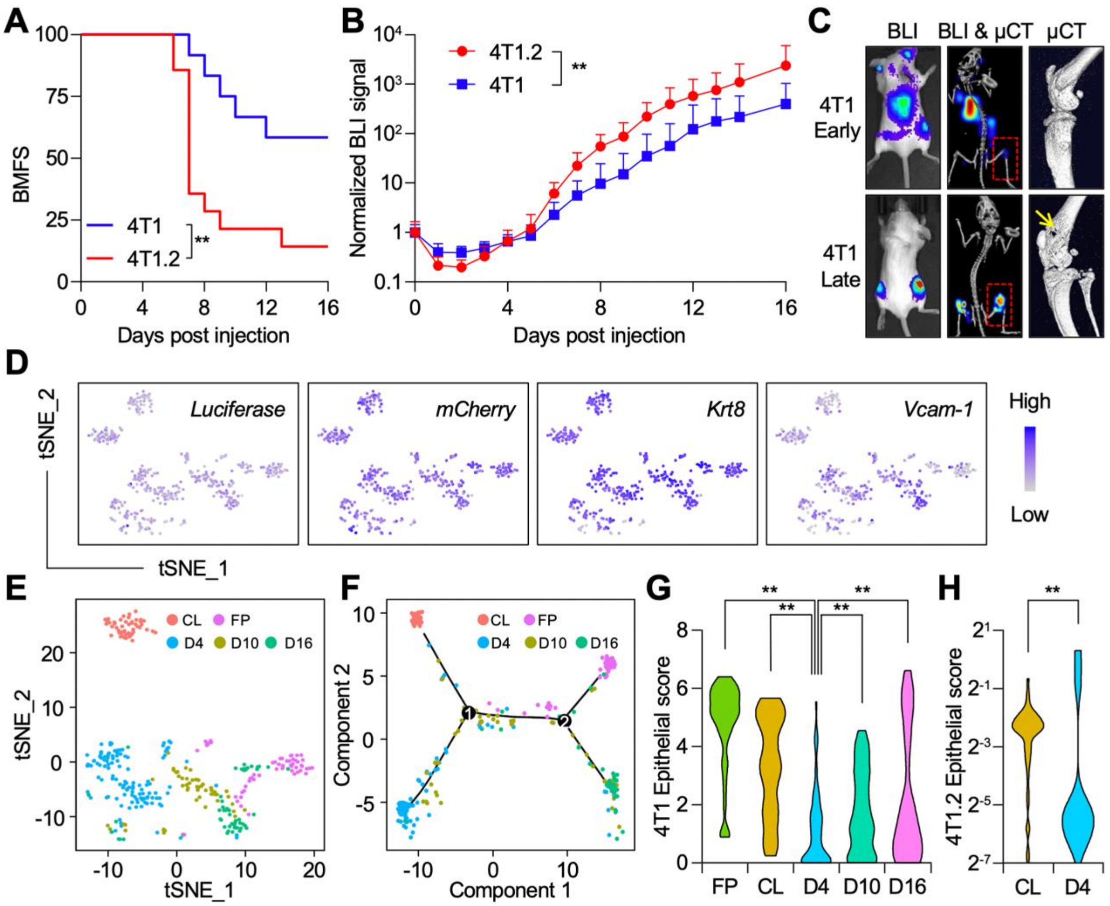


**Figure S1, related to Figure 1: Different Sources of Tumor Cells Present Distinct Gene Expression Pattens** (**A**) 4T1 or 4T1.2 cells were IC injected into 6-week-old female nude mice. Bone metastasis burden was monitored by BLI imaging. Kaplan-Meier curves for BMFS were displayed. n = 12 for 4T1 group, and n = 14 for 4T1.2 group. $^{**}p < 0.01$ by log-rank test. (**B**) Quantification of BLI signal from experiment **A**. n = 12 for 4T1 group, and n = 14 for 4T1.2 group. Data presented as mean ± SD. $^{**}p < 0.01$ by two-way repeated measures ANOVA. (**C**) Representative BLI and µCT images for early- and late-stages bone metastasis generated by 4T1 cells. Yellow arrow in µCT image points to the severely damaged osteolytic bone area. (**D**) t-SNE plots of the specific marker gene expression in the cell population. *Firefly-luciferase* and *mCherry* were exogenously labeled markers for tumor cells, *Krt8* marks epithelial cell lineage, *Vcam-1* is a previously identified bone metastatic gene. (**E**) t-SNE analysis of 4T1 cells from 4T1-CL, 4T1-FP, 4T1-D4, 4T1-D10, and 4T1-D16. Cells from different sources were color coded. (**F**) Pseudotime analysis of 4T1 cells from 4T1-CL, 4T1-FP, 4T1-D4, 4T1-D10, and 4T1-D16. Cells from different sources were color coded. (**G**) Violin plot depicts the epithelial score of 4T1 cells of 4T1-CL, 4T1-FP, 4T1-D4, 4T1-D10, and 4T1-D16 based on scRNA-seq results. **$p < 0.01$ by unpaired t-test. (**H**) Violin plot depicts the epithelial score of 4T1.2-CL and 4T1.2-D4 based on scRNA-seq results. **$p < 0.01$ by unpaired t-test.

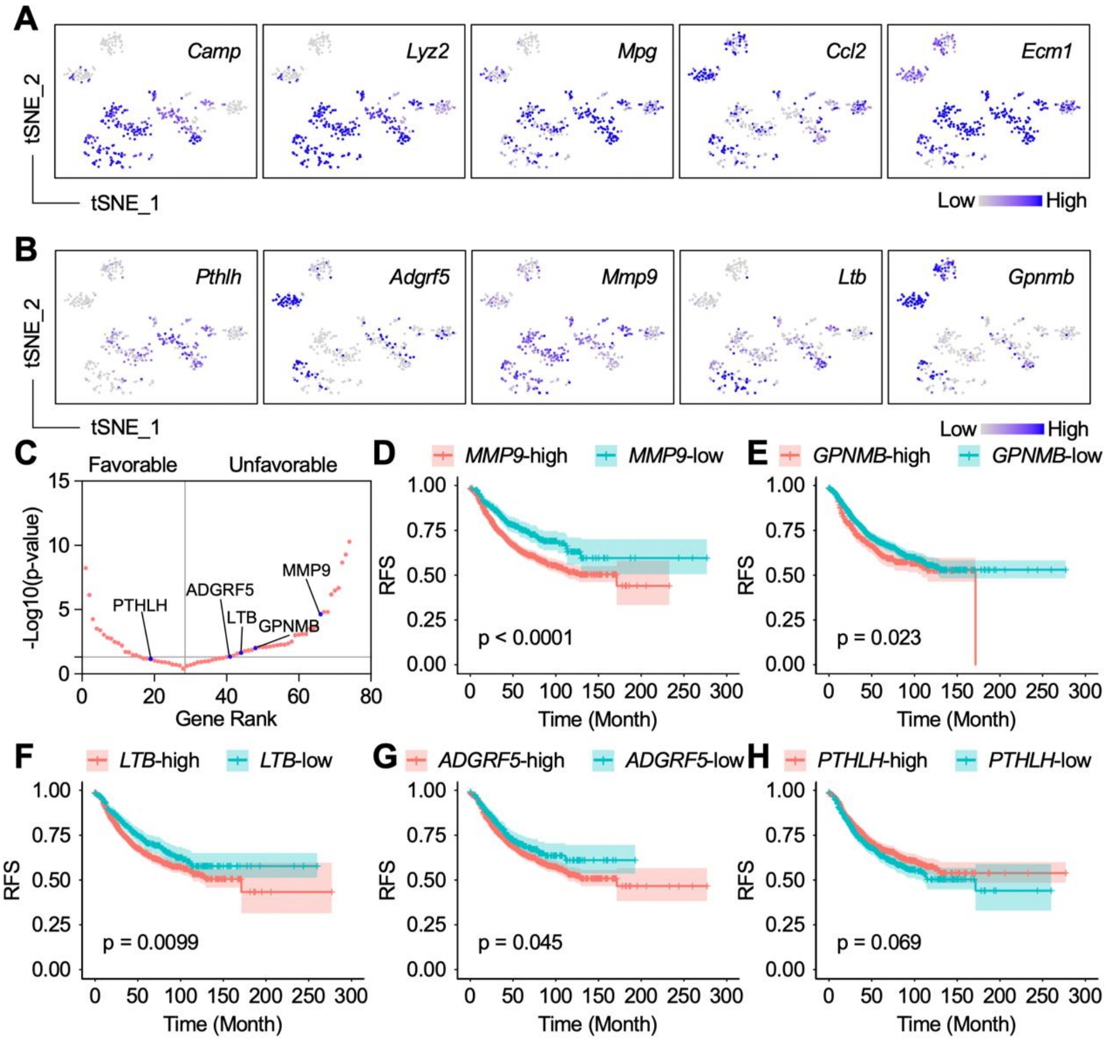


**Figure S2, related to Figure 2. Prognosis analysis of candidate genes in breast cancer patient dataset.** (**A**) By comparing 4T1-D4 *v.s.* 4T1-CL group and by comparing 4T1.2-D4 *v.s.* 4T1-D4, the genes with enriched expression were selected in a combined gene list in Supplementary Table 2. The representative gene expression patterns of the top five candidate genes were shown here in t-SNE plots. (**B**) The gene expression patterns of top five candidate genes from *in vivo* mini-cDNA library screening were shown here in t-SNE plots. (**C**) Gene ranked by the p-values from Kaplan-Meier curve of relapse-free survival in lymph node positive breast cancer patients. Patient dataset is from Kaplan-Meier Plotter database (kmplot.com), stratified by the expression of individual candidate gene. Genes on the left are those whose higher expressions are associated with better patient survival; while genes on the right are those whose higher expressions are associated with worse patient survival. (**D-H**) Kaplan-Meier plots of relapse-free survival in lymph node positive breast cancer patients, stratified by the expression of five top candidate genes (*MMP9*, *GPNMB*, *LTB*, *ADGRF5*, and *PTHLH*).

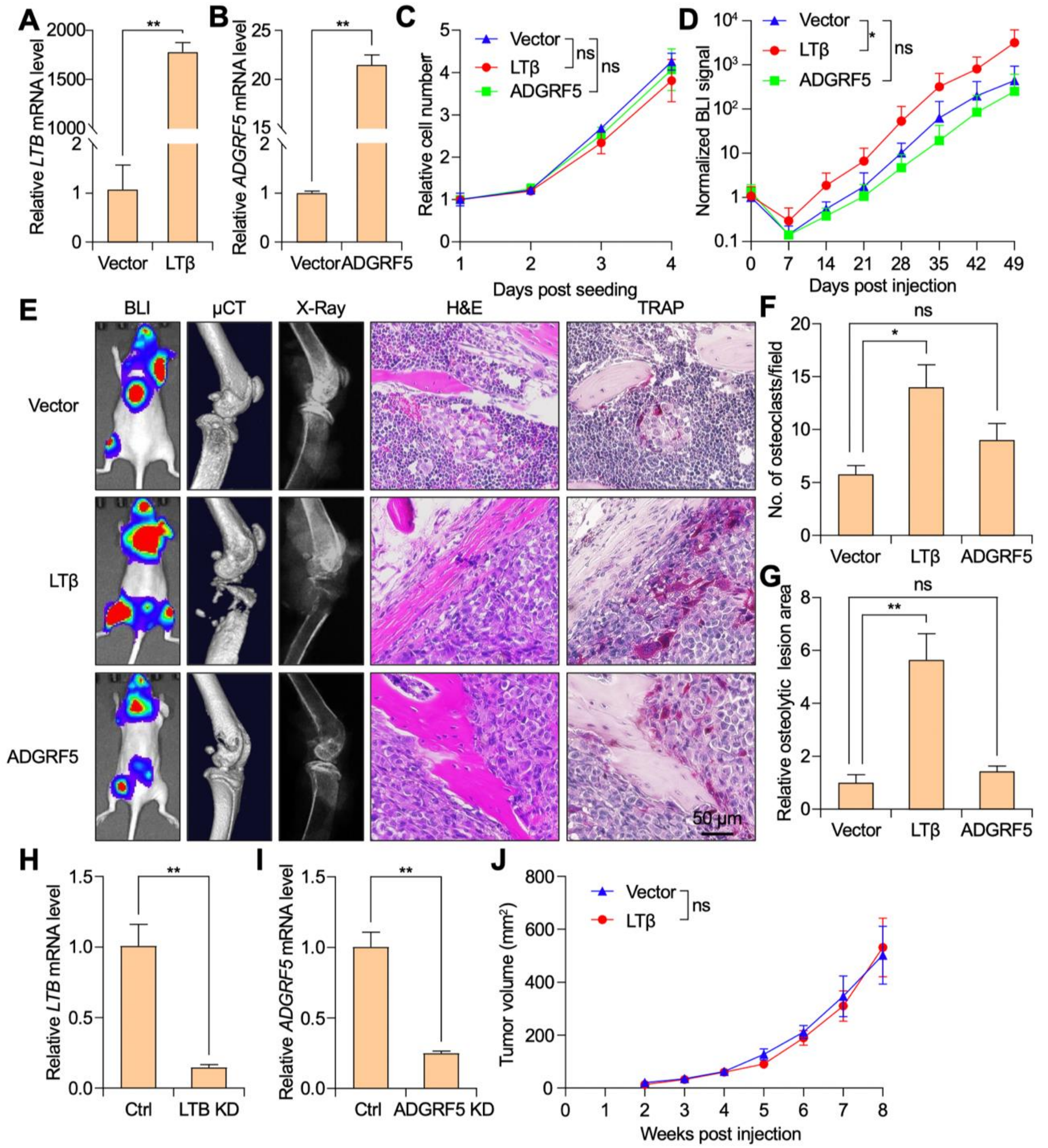


**Figure S3, related to Figure 3: LTβ Promotes Bone Metastasis.** (**A-B**) Lentivirus containing *LTB* and *ADGRF5* cDNA CDS were transduced into SCP28 cells to generate stable overexpression cell lines. The mRNA expression levels of *LTB* and *ADGRF5* in SCP28 cells were determined by qPCR, *GAPDH* was used as internal control. n = 3 per group. Data presented as mean ± SD. **p < 0.01 by unpaired t-test. (**C**) $10^3$ SCP28 cells with -Vector control, LTβ-OE, or ADGRF5-OE were seeded into 96 well plate for cell culture. Relative cell number was monitored by *in vitro* luciferase assay. n = 3 for each group. Data presented as mean ± SD. “ns” means not statistically different by two-way repeated measures ANOVA. (**D**) $10^5$ SCP28 cells with -Vector control, LTβ-OE, or ADGRF5-OE were IC injected into 6-week-old female nude mice. Bone metastasis burden was monitored by BLI imaging. n = 5 for Vector control group, n = 7 for LTβ group, and n = 7 for ADGRF5 group. Data presented as mean ± SD. ns: $p > 0.05$, *$p < 0.05$ by two-way repeated measures ANOVA. (**E**) Representative BLI, μCT, X-ray, H&E staining, and TRAP staining images of mice from experiment performed in **D**. B, bone tissue area; T, tumor area. Scale bar = 50 μm. (**F**) Quantification of the number of $TRAP^+$ osteoclasts based on TRAP staining images from experiment performed in **E**. n = 4 per group. Data presented as mean ± SEM. ns: $p > 0.05$, *$p < 0.05$ by unpaired t-test. (**G**) Quantification of osteolytic lesion areas based on X-ray images of mice from experiment performed in E. n = 5 for Vector control group, n = 7 for LTβ group, and n = 7 for ADGRF5 group. Data presented as mean ± SEM. ns: $p > 0.05$, **$p < 0.01$ by unpaired t-test. (**H-I**) LTβ and ADGRF5 in SCP28 cells were knocked down by shRNA constructs, scramble shRNA was utilized as negative control. The mRNA expression levels of these two genes were determined by qPCR, *GAPDH* was used as internal control. n = 3 per group. Data presented as mean ± SD. **$p < 0.01$ by unpaired t-test. (**J**) SCP28 cells with -Vector control or LTβ-OE were injected in the mammary fat pads of 6-week-old female nude mice. Tumor volume was measured weekly. n = 5 mice per group. Data presented as mean ± SD. ns: $p > 0.05$ by two-way repeated measures ANOVA.

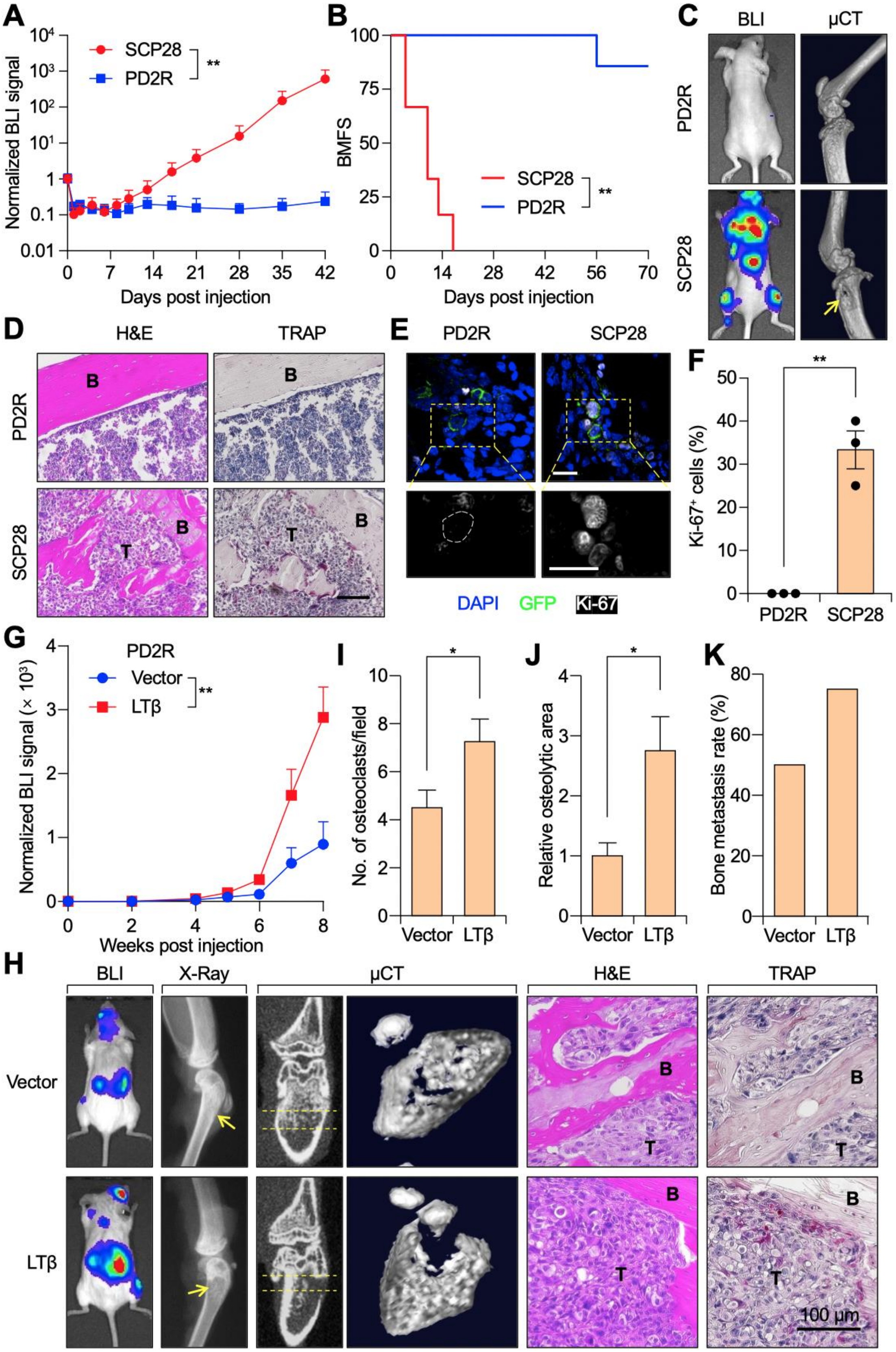

A
Normalized BLI signal
SCP28
PD2R
**
Days post injection
B
BMFS
SCP28
PD2R
**
Days post injection
C
BLI
µCT
PD2R
SCP28
D
H&E
TRAP
PD2R
SCP28
B
T
E
PD2R
SCP28
DAPI
GFP
Ki-67
F
Ki-67+ cells (%)
**
PD2R
SCP28
G
PD2R
Vector
LTβ
**
Normalized BLI signal (× 10³)
Weeks post injection
I
No. of osteoclasts/field
*
Vector
LTβ
J
Relative osteolytic area
*
Vector
LTβ
K
Bone metastasis rate (%)
Vector
LTβ
H
BLI
X-Ray
µCT
H&E
TRAP
Vector
LTβ
B
T
100 µm

**Figure S4, related to Figure 4. Pro-metastatic Activity of LTβ in Additional Bone Metastasis Models.** (**A**) SCP28 or PD2R cells were IC injected into 6-week-old female nude mice. Bone metastasis burden was monitored by BLI imaging. n = 6 for SCP28 group, and n = 7 for PD2R group. Data presented as mean ± SD. **p < 0.01 by two-way repeated measures ANOVA. (**B**) Kaplan-Meier curve of mice in experiments performed in **A**. n = 6 for SCP28 group, and n = 7 for PD2R group. **p < 0.01 by log-rank test. (**C**) Representative BLI and μCT images from mice IC injected either PD2R or SCP28 cells. Arrows indicate the osteolytic bone areas. (**D**) Representative H&E and TRAP staining images from mice IC injected either PD2R or SCP28 cells. B, bone tissue area; T, tumor area. Arrows indicate the osteolytic bone areas. Scale bar = 100 μm. (**E**) Representative IF images of Ki-67 staining in the bone tissues from PD2R or SCP28 IC-injected mice. Mice were sacrificed at Week 1 post injection for hind limb bone collection and IF staining against Ki-67. GFP: cancer cells. Scale bar = 20 μm, white dotted lines: nuclear border of $GFP^+$ cells. (**F**) Percentage of Ki-$67^+$ cells detected by IF staining from experiment in **E**. n = 3 mice for each group. Data presented as mean ± SEM. **p < 0.01 by unpaired t-test. (**G**) Similar experiment was performed as in **Fig. 4A**, expect that NSG mice were used in this experiment. Bone metastasis burden was determined weekly by BLI imaging. Quantification of BLI signal from was presented. n = 8 mice per group. **p < 0.01 by two-way repeated measures ANOVA. (**H**) Representative BLI, X-ray, μCT, H&E staining, and TRAP staining images of bone metastasis from experiment in **G**. Scale bar = 100 μm. (**I**) Quantification of the number of $TRAP^+$ osteoclasts based on TRAP staining images from experiment performed in G. n = 8 per group. *p < 0.05 by unpaired t-test. (**J**) Quantification of osteolytic lesion areas based on X-ray images of mice from experiment performed in **G**. n = 8 per group. Data presented as mean ± SEM. *p < 0.05 by unpaired t-test. (**K**) Quantification of the percentage of BLI-positive legs from *ex vivo* BLI imaging of mice from experiment in **G**. **See also Supplementary Videos 4-5.**

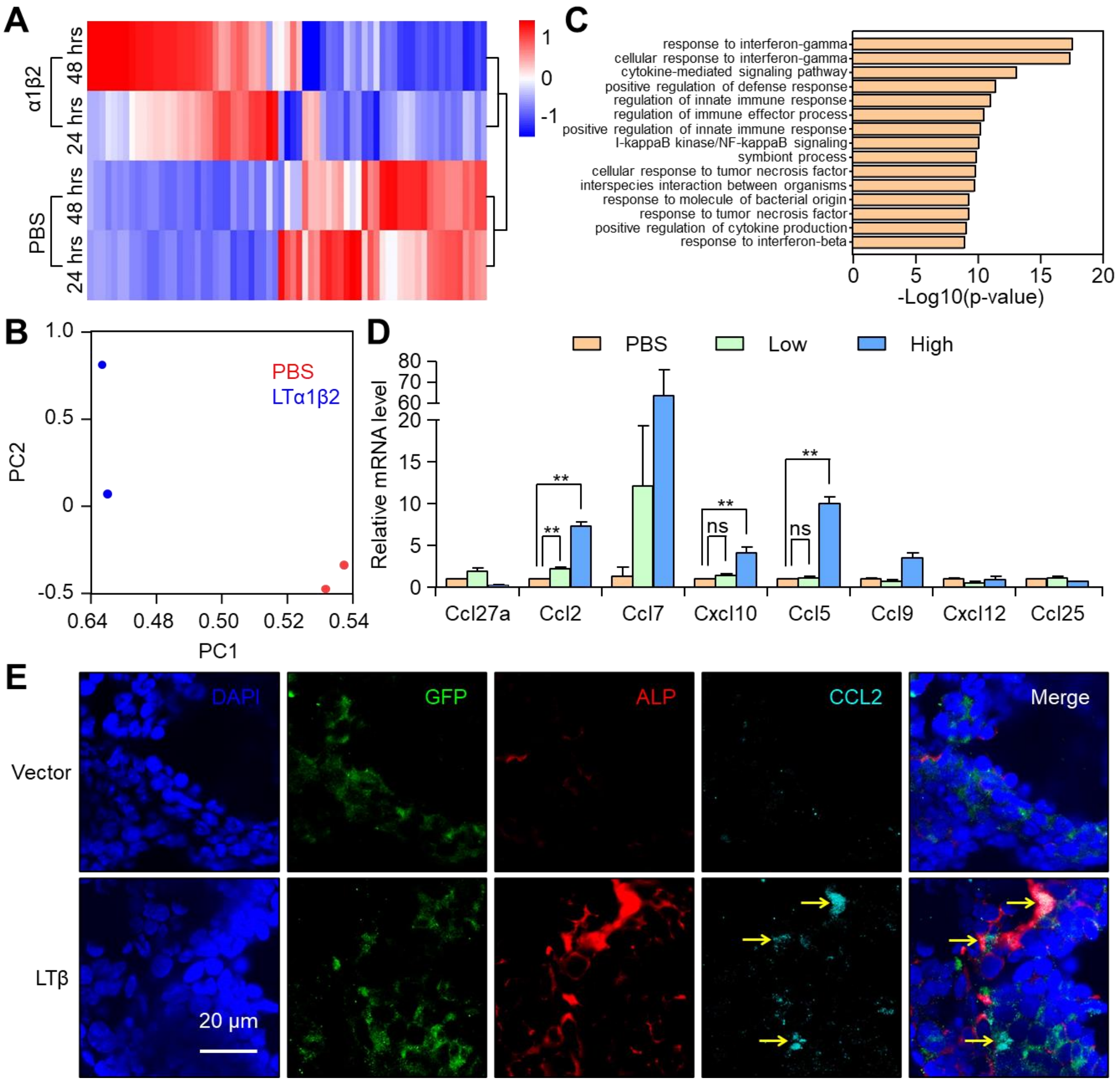


**Figure S5, related to Figure 5: Bone-seeding Tumor Cells Interact with Osteoblast Lineage Cells to Generate a Cytokine-rich Microenvironment.** (**A**) Heatmap represents differentially expressed genes in MC3T3 which were treated with either PBS or LTα1β2 for 24 and 48 hours. Scale bar indicates Z-scores. (**B**) PCA analysis demonstrated that the gene expression profiles of LTα1β2-treated MC3T3 cells were significantly different from that of PBS-treated cells. (**C**) GO analysis of enriched terms in MC3T3 treated with LTα1β2 compared to that of PBS treatment. (**D**) Relative mRNA expression levels of chemokines in MC3T3 cells treated with either PBS as control, low concentration or high concentration of recombinant LTα1β2. n = 3 per group. Data presented as mean ± SD. ns: $p > 0.05$, **$p < 0.01$ by unpaired t-test. (**E**) Representative

IF images of ALP and CCL2 staining in the bone tissues from SCP28 injected mice. Mice were sacrificed at Week 2 post injection for hind limb bone collection and IF staining against ALP and CCL2. GFP: cancer cells. ALP: osteoblasts. Yellow arrows indicate CCL2 staining. Scale bar = 20 μm.

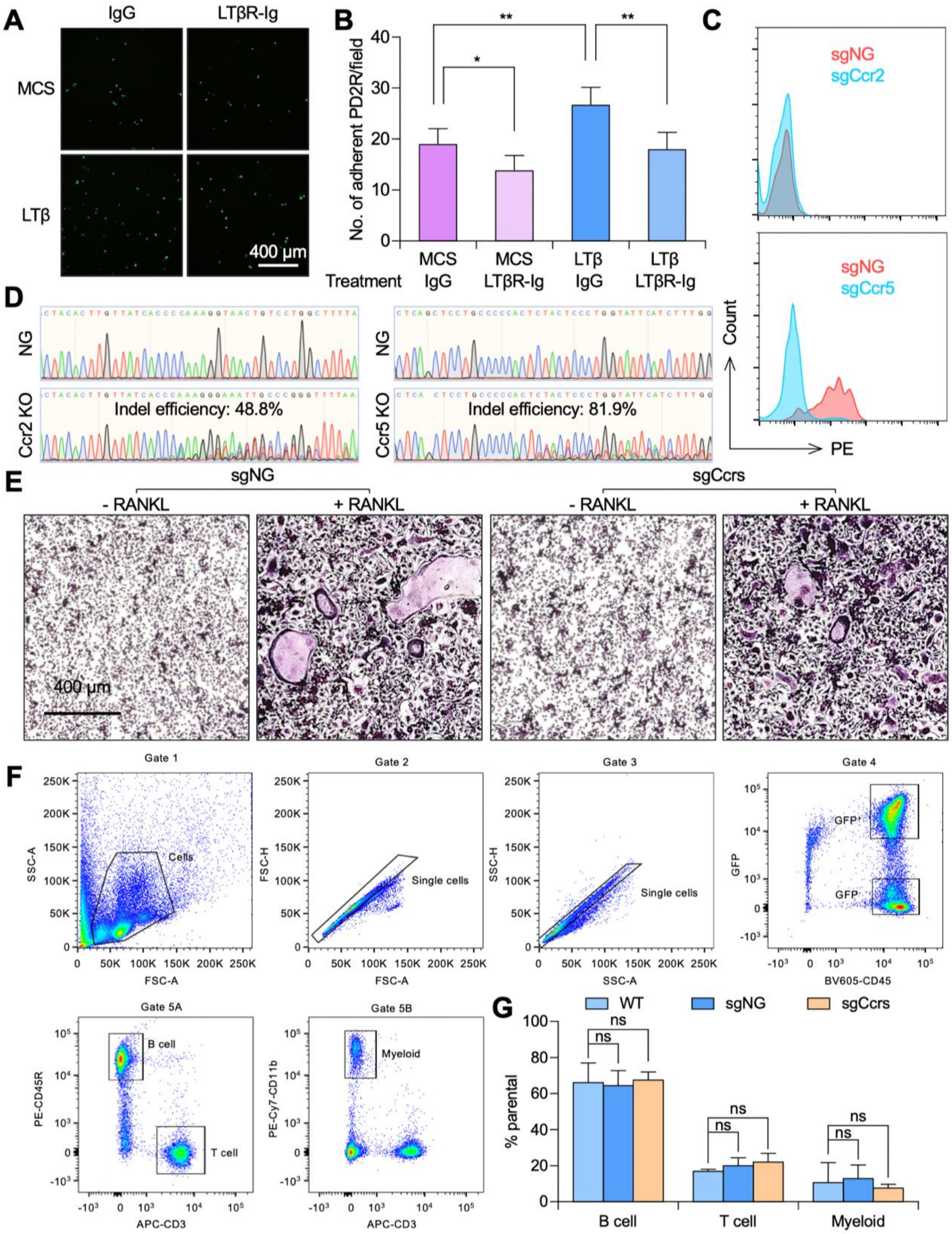


**Figure 6, related to Figure 6: CCL2/5 Promotes Osteoclastogenesis.** (**A**) Representative microscopic imaging of adhered GFP-labeled SCP28 cells was presented from experiment in **Fig. 6B**. Scale bar = 400 μm. (**B**) mCherry-labeled MC3T3 cells were seeded onto cell culture plates to reach 100% confluency. GFP-labeled PD2R cells with either -Vector or -LTβ expression were seeded on top of the MC3T3 cells for 2 minutes. The number of adhered PD2R cells was determined using direct microscopic imaging. n = 6 per group. Data presented as mean ± SD. *p < 0.05, **p < 0.01 by unpaired t-test. (**C**) Bone marrow $CD117^+$ cells from Cas9 mice were FACS analyzed for Ccr2 and Ccr5 expression a week after sgRNA transduction. Notice a clear down-regulation of Ccr5 expression after the transduction of Ccr sgRNA (sgCcr5 group). Ccr2 expression is very limited even in control group (sgNG). (**D**) Same sgRNAs against Ccr2 and Ccr5 from experiment in **C** were tested in Raw264.7 cells. DNA from these cells were then PCR and sequenced for testing the knockout efficiency. (**E**) Control BM $CD117^+$ cells or Ccr2s knockout (sgCcrs) BM $CD117^+$ cells from **Fig. 6F** were further induced for osteoclast differentiation *in vitro*. Osteoclast cells were visualized by an osteoclast-staining kit. (**F**) The peripheral blood of transplanted recipient mice was collected and analyzed by FACS analysis, using the gating schemes as shown in the figure panels. FSC-A, forward scatter area; SSC-A, side scatter area; FSC-H, forward scatter height; SSC-H, side scatter height. For transplanted recipient mice: $GFP^+$ cells, which were differentiated from successfully transfected BM $CD117^+$ cells, were used for further analysis of the percentage of B cells, T cells, and myeloid cells. For wild type mice without transplantation, $GFP^-$ cells were similarly analyzed as negative control. (**G**) Quantification of the percentages of B cells, T cells, and myeloid cells from mice without or with transplantation. n = 5 mice for each group. Data presented as mean ± SEM. ns: p > 0.05 by unpaired t-test.

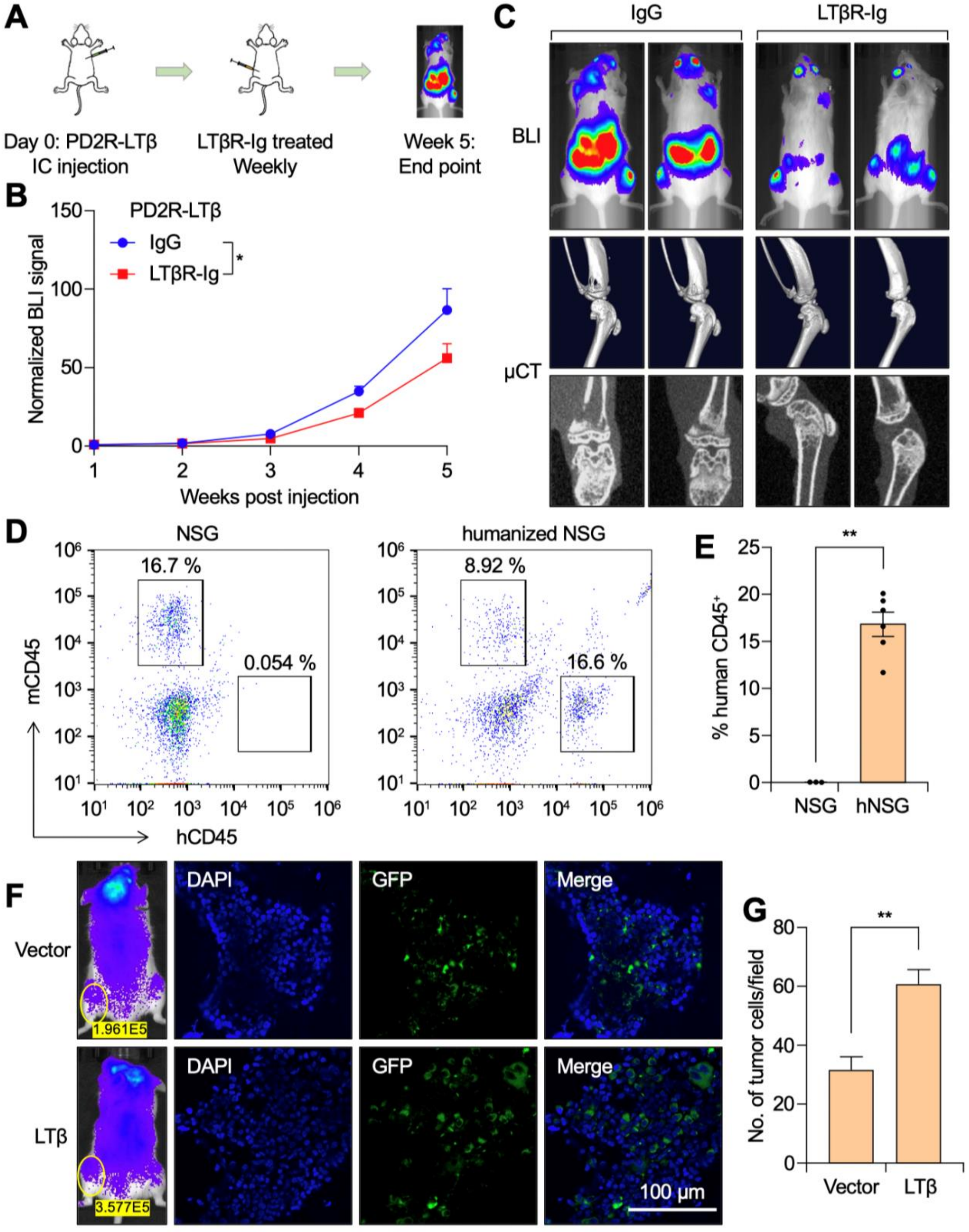


**Figure S7, related to Figure 7: LTβ Mediated Signaling is Essential to Bone Metastasis Progression.** (**A**) PD2R-LTβ cells were IC injected into 6-week-old female NSG mice. Mice were treated with LTβR-Ig recombinant protein three days later and continued until the experimental endpoint. (**B**) Bone metastasis burden was monitored by BLI imaging from experiment in **A**. n = 9 mice per each group. Data presented as mean ± SD. *$p < 0.05$ by two-way repeated measures ANOVA. (**C**) Representative BLI and μCT images of bone metastasis from experiment in **A**. (**D**) Humanized mouse model: human HSC cells were inoculated into lethally irradiated NSG mice to reconstitute their hematopoietic system. Peripheral blood from these mice were FACS analyzed for the presence of human $CD45^+$ cells. Representative FACS-plot from this experiment was shown. (**E**) Quantification of the percentage of human $CD45^+$ cells from control NSG mice and from humanized mice. n = 3 for control group and n = 6 for humanized mouse group. Data presented as mean ± SD. **$p < 0.01$ by unpaired t-test. (**F**) GFP-labeled SCP28 cells with Vector control or LTβ OE were IC injected into humanized NSG mice. Mice were sacrificed 5 days later to collect hindlimbs for IF staining. Samples were counter-stained with DAPI. Scale bar = 100 μm. (**G**) Quantification of $GFP^+$ cells per field from experiment in **F**. n = 8 per group. Data presented as mean ± SEM. **$p < 0.01$ by unpaired t-test.
