## Supplemental Table 1 for "Lymphotoxin*-*β Promotes Bone Colonization and Osteolytic Outgrowth of Indolent Bone Metastatic cells of Breast Cancer"

**Table S1. The correlation between cell types and clusters in t-SNE plot**

| Cluster | 4T1-FP | 4T1-CL | 4T1-D4 | 4T1-D10 | 4T1-D16 | 4T1.2-CL | 4T1.2-D4 | Total |
| --- | --- | --- | --- | --- | --- | --- | --- | --- |
| 0 | 6 | 0 | 12 | 35 | 4 | 0 | 0 | 57 |
| 1 | 1 | 0 | 46 | 2 | 1 | 0 | 0 | 50 |
| 2 | 0 | 0 | 0 | 0 | 0 | 49 | 0 | 49 |
| 3 | 0 | 48 | 0 | 0 | 0 | 0 | 0 | 48 |
| 4 | 0 | 0 | 46 | 1 | 0 | 0 | 0 | 47 |
| 5 | 1 | 0 | 11 | 9 | 7 | 0 | 17 | 45 |
| 6 | 3 | 0 | 2 | 5 | 27 | 0 | 0 | 37 |
| 7 | 34 | 0 | 0 | 0 | 2 | 0 | 0 | 36 |
| 8 | 0 | 0 | 1 | 0 | 0 | 0 | 31 | 32 |
| 9 | 11 | 0 | 1 | 0 | 11 | 0 | 0 | 23 |
| Total | 56 | 48 | 119 | 52 | 52 | 49 | 48 | 424 |
