## Supplemental Table 2 for "Lymphotoxin*-*β Promotes Bone Colonization and Osteolytic Outgrowth of Indolent Bone Metastatic cells of Breast Cancer"

**Table S2. Candidates for cDNA library construction.**

| **Symble** | **Gene** | **avg_logFC** | **p_val** | **From *** | **Location** |
| --- | --- | --- | --- | --- | --- |
| Camp | cathelicidin antimicrobial peptide | 7.251272356 | 6.42583E-23 | A | Secreted |
| Lyz2 | lysozyme | 5.705667086 | 2.52464E-23 | A | Secreted |
| Mgp | N-methylpurine DNA glycosylase | 4.419853756 | 9.77163E-17 | A | Secreted |
| Ccl2 | C-C motif chemokine ligand 2 | 3.63426302 | 6.44425E-12 | B | Secreted |
| Ecm1 | extracellular matrix protein 1 | 3.270627567 | 3.43673E-22 | A | Secreted |
| Selp | selectin P | 3.202171034 | 9.12776E-14 | A | Membrane |
| Aqp1 | aquaporin 1 | 3.183380058 | 3.80408E-21 | A | Membrane |
| Tmem41a | transmembrane protein 41A | 3.137487161 | 9.4308E-16 | B | Membrane |
| Plau | plasminogen activator, urokinase | 3.025414992 | 9.93258E-10 | B | Secreted |
| Tyrobp | transmembrane immune signaling adaptor TYROBP | 2.950529121 | 2.63543E-23 | A | Membrane |
| Kazald1 | Kazal type serine peptidase inhibitor domain 1 | 2.854350819 | 1.14891E-09 | A | Secreted |
| Nptx1 | neuronal pentraxin 1 | 2.831248284 | 6.16396E-12 | A | Membrane |
| Adrb2 | adrenoceptor beta 2 | 2.798134965 | 1.82925E-17 | A | Membrane |
| Ltf | lactotransferrin | 2.78888383 | 2.52464E-23 | A | Membrane |
| Mmp3 | matrix metallopeptidase 3 | 2.758579113 | 1.63004E-07 | A | Secreted |
| Pthlh | parathyroid hormone like hormone | 2.724906774 | 4.00628E-08 | A | Secreted |
| Dmp1 | dentin matrix acidic phosphoprotein 1 | 2.495273389 | 3.07774E-07 | A | Secreted |
| Ptprcap | protein tyrosine phosphatase receptor type C associated protein | 2.491285506 | 9.26343E-10 | B | Membrane |
| Slc2a5 | solute carrier family 2 member 5 | 2.46786836 | 1.88941E-25 | B | Membrane |
| Pla2g7 | phospholipase A2 group VII | 2.450041432 | 3.73946E-18 | A | Secreted |
| Cx3cl1 | C-X3-C motif chemokine ligand 1 | 2.345369551 | 1.70341E-16 | A | Membrane |
| Icam2 | intercellular adhesion molecule 2 | 2.335802857 | 4.87511E-10 | B | Membrane |
| Col6a1 | collagen type VI alpha 1 chain | 2.323587298 | 1.19999E-15 | A | Secreted |
| Mmp9 | matrix metallopeptidase 9 | 2.315937457 | 1.22587E-15 | A | Secreted |
| Il2rg | interleukin 2 receptor subunit gamma | 2.096477058 | 6.04893E-08 | B | Membrane |
| Cd82 | CD82 molecule | 2.08113289 | 1.61484E-17 | A | Membrane |
| Ttyh2 | tweety family member 2 | 2.05310898 | 1.85921E-10 | A | Membrane |
| Cldn9 | claudin 9 | 2.041983567 | 4.29583E-10 | B | Membrane |
| Serpinf1 | serpin family F member 1 | 2.02495171 | 7.61442E-08 | B | Secreted |
| Cd177 | CD177 molecule | 2.007790157 | 5.58926E-23 | A | Membrane |
| Wnt6 | Wnt family member 6 | 1.98189538 | 1.96599E-09 | B | Secreted |
| Plvap | plasmalemma vesicle associated protein | 1.937845879 | 1.39382E-11 | B | Membrane |
| Vasn | vasorin | 1.900669311 | 6.97657E-09 | A | Membrane |
| Lcn2 | lipocalin 2 | 1.882831905 | 3.38623E-15 | A | Secreted |
| Ecscr | endothelial cell surface expressed chemotaxis and apoptosis regulator | 1.853773049 | 5.06748E-07 | A | Membrane |
| Loxl3 | lysyl oxidase like 3 | 1.844912919 | 2.16217E-11 | A | Secreted |
| Angptl2 | angiopoietin like 2 | 1.84307706 | 4.08807E-19 | A | Secreted |
| Pcolce | procollagen C-endopeptidase enhancer | 1.82958385 | 2.6754E-15 | A | Secreted |
| Atp1b1 | ATPase Na+/K+ transporting subunit beta 1 | 1.804461401 | 2.31163E-08 | B | Membrane |
| Comt | catechol-O-methyltransferase | 1.794276438 | 9.75043E-16 | B | Membrane |
| Itgb7 | integrin subunit beta 7 | 1.763017937 | 1.70686E-08 | B | Membrane |
| Adgrg1 | adhesion G protein-coupled receptor G1 | 1.744132038 | 6.16621E-13 | A | Membrane |
| Ngfr | nerve growth factor receptor | 1.732572054 | 6.19538E-13 | B | Membrane |
| Spp1 | secreted phosphoprotein 1 | 1.653905784 | 8.03098E-12 | A | Secreted |
| Ctss | cathepsin S | 1.648549965 | 5.58926E-23 | A | Secreted |
| Tnfrsf19 | TNF receptor superfamily member 19 | 1.638336455 | 1.19463E-10 | B | Membrane |
| Hp | haptoglobin | 1.622391251 | 1.50127E-22 | A | Secreted |
| Flrt3 | fibronectin leucine rich transmembrane protein 3 | 1.570183415 | 7.59135E-09 | A | Membrane |
| Pmp22 | peripheral myelin protein 22 | 1.557469067 | 9.15858E-09 | A | Membrane |
| Slc5a3 | solute carrier family 5 member 3 | 1.545143019 | 1.08634E-14 | A | Membrane |
| Mcemp1 | mast cell expressed membrane protein 1 | 1.507669512 | 1.37322E-22 | A | Membrane |
| Efnb1 | ephrin B1 | 1.496717151 | 5.96598E-09 | A | Membrane |
| Ltb | lymphotoxin beta | 1.4945106 | 8.44571E-07 | B | Membrane |
| Srgn | serglycin | 1.457866535 | 2.25002E-23 | A | Secreted |
| Slc29a1 | solute carrier family 29 member 1 | 1.41400065 | 1.07743E-16 | A | Membrane |
| Csf1 | colony stimulating factor 1 | 1.403221774 | 8.94514E-08 | B | Membrane |
| Sdc2 | syndecan 2 | 1.400451288 | 6.61374E-16 | A | Membrane |
| Tfpi | tissue factor pathway inhibitor | 1.392287519 | 4.11989E-15 | B | Secreted |
| Tgfa | transforming growth factor alpha | 1.384637313 | 2.52598E-12 | A | Membrane |
| Lrp11 | LDL receptor related protein 11 | 1.337398667 | 1.28752E-07 | B | Membrane |
| Vcam1 | vascular cell adhesion molecule 1 | 1.336338989 | 5.97904E-10 | A | Membrane |
| Tinagl1 | tubulointerstitial nephritis antigen like 1 | 1.321060758 | 2.87421E-13 | A | Secreted |
| Ier3 | immediate early response 3 | 1.291405198 | 1.63055E-10 | A | Membrane |
| Fbln1 | fibulin 1 | 1.279918476 | 1.98555E-13 | B | Secreted |
| Eva1c | eva-1 homolog C | 1.243573062 | 1.23898E-07 | A | Membrane |
| Tgfbi | transforming growth factor beta induced | 1.24306669 | 1.09417E-18 | A | Secreted |
| Ccl4 | C-C motif chemokine ligand 4 | 1.225418045 | 1.32986E-17 | A | Secreted |
| Sspn | sarcospan | 1.221901021 | 1.28011E-07 | A | Membrane |
| Pglyrp1 | peptidoglycan recognition protein 1 | 1.219590753 | 5.73063E-20 | A | Secreted |
| Ghr | growth hormone receptor | 1.182412198 | 5.23178E-10 | A | Membrane |
| Fgfr2 | fibroblast growth factor receptor 2 | 1.165764588 | 8.48343E-13 | A | Membrane |
| Siglecg | sialic acid binding Ig like lectin 10 | 1.151150221 | 9.0055E-10 | B | Membrane |
| Clcn2 | chloride voltage-gated channel 2 | 1.13973782 | 4.75395E-07 | B | Membrane |
| Arvcf | ARVCF delta catenin family member | 1.115218532 | 2.34087E-08 | B | Membrane |
| Afp | alpha fetoprotein | 1.108704335 | 2.78914E-08 | B | Secreted |
| Tfrc | transferrin receptor | 1.086732424 | 1.48638E-07 | B | Membrane |
| Gpnmb | glycoprotein nmb | 1.07863208 | 1.01894E-11 | B | Membrane |
| Cd37 | CD37 molecule | 1.046604656 | 4.65315E-10 | B | Membrane |
| Cd69 | CD69 molecule | 1.043927144 | 1.9914E-10 | B | Membrane |
| Cpa3 | carboxypeptidase A3 | 1.04375105 | 2.03419E-07 | B | Secreted |
| Ap2m1 | adaptor related protein complex 2 subunit mu 1 | 1.042390939 | 1.9139E-08 | B | Membrane |
| Pcdh1 | protocadherin 1 | 1.030079388 | 3.24338E-08 | B | Membrane |
| Adgrf5 | adhesion G protein-coupled receptor F5 | 0.970358078 | 1.39049E-17 | B | Membrane |
| Cd63 | CD63 molecule | 0.634745506 | 5.50437E-08 | B | Membrane |

Note:

* **A**, 4T1 cells from Day 4 post injection versus 4T1 cell line; **B**, 4T1.2 cells from Day 4 post injection versus 4T1 cells from day 4 post injection.
