## Supplemental Table 3 for "Lymphotoxin*-*β Promotes Bone Colonization and Osteolytic Outgrowth of Indolent Bone Metastatic cells of Breast Cancer"

**Table S3. DNA Oligos for plasmid construction. Related to STAR Methods.**

| **Primer** | **Sequence (5’-3’)** |
| --- | --- |
| FLAG-10N-XhoI-F | AGCATCTAGAGCGGCCGCCGATTACAAGGATGACGACGATAAGTGATGCAGGCTACNNNNNNNNNNGGGTAGGGGAGGCGCTTTTCCCAA |
| MPGK-AgeI-R | GTTGGCGCCTACCGGTGGATGTGGA |
| mCherry-XhoI-F | GGAATTAGATCTCTCGAGGTTAACGAATTCGCCACCATGGTGAGCAAGGGCGAGGAGGAT |
| mCherry-SalI-R | CTGCAGGTCGACCTACTTGTACAGCTCGTCCATGCC |
