## Supplemental Table 4 for "Lymphotoxin*-*β Promotes Bone Colonization and Osteolytic Outgrowth of Indolent Bone Metastatic cells of Breast Cancer"

**Table S4. Oligos for NGS library construction. Related to STAR Methods.**

| **Primer** | **Sequence (5’-3’)** |
| --- | --- |
| NGS-F | CTTTCCCTACACGACGCTCTTCCGATCTGATTACAAGGATGACGACGATAAG |
| NGS-R | GTGACTGGAGTTCAGACGTGTGCTCTTCCGATCTGGCCTACCCGCTTCCATTGCTCAG |
| NGS-Universal-F | AATGATACGGCGACCACCGAGATCTACACTCTTTCCCTACACGACGCTCT |
| NGS-Index-R | CAAGCAGAAGACGGCATACGAGAT-index-GTGACTGGAGTTCAGACGTG |
