## Supplemental Table 5 for "Lymphotoxin*-*β Promotes Bone Colonization and Osteolytic Outgrowth of Indolent Bone Metastatic cells of Breast Cancer"

**Table S5. shRNA sequences. Related to STAR Methods.**

| **shRNA** | **Sequence (5’-3’)** |
| --- | --- |
| Scramble shRNA | CCGGCCTAAGGTTAAGTCGCCCTCGCTCGAGCGAGGGCGACTTAACCTTAGGTTTTTG |
| LTB shRNA | CCGGGCGAGAGGGAAGACCTTCTTTCTCGAGAAAGAAGGTCTTCCCTCTCGCTTTTTG |
| ADGRF5 shRNA | CCGGGCCCAACTTCTCTGTCTATATCTCGAGATATAGACAGAGAAGTTGGGCTTTTT |
