## Supplemental Table 6 for "Lymphotoxin*-*β Promotes Bone Colonization and Osteolytic Outgrowth of Indolent Bone Metastatic cells of Breast Cancer"

**Table S6. Oligos for qPCR. Related to STAR Methods.**

| **Gene** | **Forward (5’-3’)** | **Reverse (5’-3’)** |
| --- | --- | --- |
| *Ltb* | TGGCAGGAGCTACTTCCCT | TCCAGTCTTTTCTGAGCCTGT |
| *Ccl27a* | AGGAGGATTGTCCACATGGAA | CTTGGCGTTCTAACCACCGA |
| *Ccl2* | TTAAAAACCTGGATCGGAACCAA | GCATTAGCTTCAGATTTACGGGT |
| *Ccl7* | GCTGCTTTCAGCATCCAAGTG | CCAGGGACACCGACTACTG |
| *Cxcl10* | CCAAGTGCTGCCGTCATTTTC | GGCTCGCAGGGATGATTTCAA |
| *Ccl5* | GCTGCTTTGCCTACCTCTCC | TCGAGTGACAAACACGACTGC |
| *Ccl9* | CCCTCTCCTTCCTCATTCTTACA | AGTCTTGAAAGCCCATGTGAAA |
| *Cxcl12* | TGCATCAGTGACGGTAAACCA | TTCTTCAGCCGTGCAACAATC |
| *Ccl25* | TTACCAGCACAGGATCAAATGG | CGGAAGTAGAATCTCACAGCAC |
| *Gapdh* | AGGTCGGTGTGAACGGATTTG | TGTAGACCATGTAGTTGAGGTCA |
| *LTB* | GTACGGGCCTCTCTGGTACA | GTCCACCATATCGGGGTGAC |
| *ADGRF5* | CGGAAGGGTTACGGAATTTTACC | GTGATGGTGGTGTAGTCTTGAC |
| *GAPDH* | GAAGGTGAAGGTCGGAGTC | GAAGATGGTGATGGGATTTC |
